# Histone H3 tail charge patterns govern nucleosome condensate formation and dynamics

**DOI:** 10.1101/2025.04.09.647968

**Authors:** Erin F. Hammonds, Anurag Singh, Krishna K. Suresh, Sean Yang, Sarah S. Meidl Zahorodny, Ritika Gupta, Davit A. Potoyan, Priya R. Banerjee, Emma A. Morrison

**Affiliations:** Department of Biochemistry, Medical College of Wisconsin, 8701 Watertown Plank Rd., Milwaukee, WI 53226; Department of Physics, The State University of New York at Buffalo, Buffalo, NY 14260; Department of Chemistry, Iowa State University, Ames, IA 50011

## Abstract

Emerging models of nuclear organization suggest that chromatin forms functionally distinct microenvironments through phase separation. As chromatin architecture is organized at the level of the nucleosome and regulated by histone post-translational modifications, we investigated how these known regulatory mechanisms influence nucleosome phase behavior. By systematically altering charge distribution within the H3 tail, we found that the terminal and central regions modulate the phase boundary and tune nucleosome condensate viscosity differentially, as revealed by microscopy-based assays, microrheology, and simulations. Nuclear magnetic resonance relaxation experiments revealed that H3 tails remain dynamically mobile within condensates, and their mobility correlates with condensate viscosity. These results demonstrate that the number, identity, and spatial arrangement of basic residues in the H3 tail critically regulate nucleosome phase separation. Our findings support a model in which nucleosomes, through their intrinsic properties and modifications, actively shape the local chromatin microenvironment—providing new insight into the histone language in chromatin condensates.

## INTRODUCTION

Dynamic chromatin organization is being re-examined through the lens of phase separation (1–6). Current models invoke phase separation and biomolecular condensates in compartmentalization and the formation of localized environments that are functionally distinct within the nucleus, which dynamically change in response to stimuli (2, 3, 5–11). Chromatin regions ranging from transcriptionally silenced heterochromatin to transcriptionally active super-enhancers form phase-separated regions (12, 13), and condensates with such diverse functions have varied physicochemical and material properties (9, 14, 15). It is appealing to consider a model wherein the nucleus exists as a collection of phase-separated regions, each enriched in a distinct set of chromatin, epigenetic modifications, proteins, and RNA that determine the local environment and inform the biological activity within.

The nucleus is rich in multivalent biopolymers capable of undergoing phase separation, and chromatin is no exception with its histone and DNA components. Chromatin can undergo phase separation in the absence of chromatin-interacting proteins, and this holds true even down to the level of the nucleosome core particle (NCP), the basic subunit of chromatin (7, 16–21). Recent reports have also shown the effect of nucleosome spacing (length of linker DNA) on nucleosome mobility and thermodynamic stability of chromatin condensates (22, 23). Within the NCP, ~147-bp DNA wraps around a folded histone core, and the intrinsically disordered termini, or tails, of the histones protrude. The histone tails are enriched in basic residues—the histone tails range from 11-36 residues in length and are 31-42% lysines and arginines while only 0-8% aspartates and glutamates. The histone tails form both intra- and inter-nucleosomal interactions with DNA within the context of nucleosomes and nucleosome arrays, and the H3 and H4 tails dominate inter-nucleosomal interactions (24–33). Chromatin phase separation is driven, at least in part, by inter-nucleosomal interactions between these highly basic histone tails and DNA (7, 16, 21). In support, removal of the H3 or H4 tails individually or all histone tails simultaneously abrogates phase separation (7, 16). Furthermore, neutralizing mutations of the H4 tail basic patch abrogate phase separation while neutralization of the H2A/H2B acidic patch does not, supporting the importance of histone tail-DNA interactions over histone tail-histone core interactions in phase separation (7). Here, we hypothesize that site-specific mutations of basic residues within histone tails alter histone–DNA electrostatic interactions, thereby modulating the condensate properties of NCPs. Because the emergent properties of chromatin subcompartments arise from the underlying physical characteristics of their constituent chromatin units, it is essential to elucidate how such molecular-level perturbations influence the material properties of nucleosome condensates.

Post-translational modifications (PTMs) on histone proteins are epigenomic marks that regulate chromatin organization, including via phase separation (34). Although histone PTMs can regulate chromatin condensates indirectly through interactions with proteins including HP1α, linker histone, and BRD4, they also directly regulate chromatin (7, 35, 36). Charge-modulating PTMs of histone tail residues, including acylation of lysine and citrullination of arginine, alter chromatin compaction and are poised to disrupt the multivalent electrostatic interactions between histone tails and DNA that drive nucleosome phase separation (37, 38). In initial investigations into the direct effects of histone PTMs, non-specific hyperacetylation of histones by p300 led to the dissolution of nucleosome array condensates (7). While both lysine and arginine carry a single positive charge at neutral pH, they differ in their interactions with DNA. Arginines are generally thought to form stronger interactions with DNA due to more favorable enthalpic contributions with the phosphate backbone, a greater hydrogen-bonding capacity, a greater propensity for cation-π interactions, and a lower energetic cost of desolvation (39–41). Greater compaction and condensation are observed with arginine-rich than lysine-rich peptides (42–47). The more favorable interaction of arginines with DNA is also observed in simulations of NCPs and nucleosomes (32, 48, 49). What remains unclear is how the position- specific contributions of lysine and arginine residues within histone tails, and their neutralization by PTMs, influence nucleosome condensate formation and dynamics.

Here, we begin to systematically dissect the influence of lysines and arginines and their charge-neutralizing modifications by investigating the role of basic residues of the H3 tail, the longest of the core histone tails (24), on nucleosome phase separation. In the case of the H3 tail, six lysines and four arginines are distributed along its length with the potential to contribute to phase separation via inter-nucleosomal interactions with DNA. We investigate the position-dependent effect of perturbing lysines or arginines in the terminal or central regions on nucleosome phase separation. Basic residues are perturbed in these regions via neutralizing mutations to glutamine as mimetics for acetylation and citrullination, or by converting all basic residues to a single type (i.e., either lysines or arginines). Using microscopy-based techniques in conjunction with simulations, we demonstrate that the number and position of lysine and arginine neutralizations within the H3 tail determine nucleosome phase boundaries and the viscosity of their condensates. NMR dynamics experiments show minimal differences in picosecond-to-nanosecond timescale dynamics of the H3 tail within condensates and display a correlation between condensate viscosity and H3 tail dynamics. Together, these data contribute to a model wherein the intrinsic properties of nucleosomes and their condensates are involved in defining the local chromatin microenvironment, which can be tuned via position-specific charge-modulating histone PTMs.

## MATERIALS AND METHODS

### Expression and purification of full-length histones

Full-length human histone sequences were expressed and purified as previously described (16, 50). These protein sequences correspond to UniProt sequences P0C0S8, P62807, Q71DI3 (with C110A), and P62805. Briefly, each histone sequence was expressed from pET3a vectors using T7 Express lysY competent *E. coli* (New England Biolabs) or Novagen Rosetta 2(DE3) pLysS competent *E. coli* (Millipore Sigma). Mutants were generated via Q5 Site-Directed Mutagenesis Kit (New England Biolabs). The histone H3 constructs used in this study were: unmodified, R2Q, R8Q, R2/8Q (2xR-Q_terminal_), R17/26Q (2xR-Q_central_), R2/8/17/26Q (4xR-Q), K4/9Q (2xK-Q_terminal_), K14/18/23/27Q (4xK-Q_central_), K4/9/18/27Q (4xK-Q_even_), K4/9/14/18/23/27Q (6xK-Q), R2/8/17/26K (All-K), K4/9/14/18/23/27R (All-R), and tailless [here, referring to H3(28-135)]. Briefly, histones were extracted from inclusion bodies and purified in Urea Buffer (7 M urea, 20 mM sodium acetate pH 5.2, 100 mM NaCl, 5 mM β-ME, 1 mM EDTA) sequentially over Q-Sepharose Fast Flow resin (GE Healthcare) and SP-Sepharose Fast Flow resin (GE Healthcare) or Proteindex IEX-CM Agarose 6 Fast Flow resin (Marvelgent Biosciences). Pure histones were dialyzed into ddH_2_O with 1 mM β-ME and stored lyophilized at −20°C until needed for nucleosome reconstitution. ESI mass spectrometry (Finnigan LTQ, Thermo Electron Corporation, San Jose, CA) was used for histone quality assessment based on molecular weight and used to ensure no unintended modifications, such as carbamylation.

### Amplification and purification of 147 bp Widom 601 DNA

DNA used in both NCP reconstitution and peptide-DNA assays was obtained using a plasmid with 32 repeats of the 147-bp Widom 601 sequence (ATCGAGAATC CCGGTGCCGA GGCCGCTCAA TTGGTCGTAG ACAGCTCTAG CACCGCTTAA ACGCACGTAC GCGCTGTCCC CCGCGTTTTA ACCGCCAAGG GGATTACTCC CTAGTCTCCA GGCACGTGTC AGATATATAC ATCCGAT). Bacterial amplification and purification of 147-bp DNA was performed as previously described (16, 50). Following transformation using NEB 5-alpha Competent *E. coli* (New England Biolabs), the bacteria were grown in 2xYT media pH 7.5. The plasmid was purified using an alkaline lysis method and phenol-chloroform extractions. The 147-bp fragments were released via digestion by EcoRV-HF enzyme (New England Biolabs) and purified from the vector using PEG precipitation and HiTrap DEAE Fast Flow Columns (GE Healthcare) via FPLC (Bio-Rad). DNA purity was assessed using 5% native-PAGE gel electrophoresis. The 147-bp DNA fragments were stored following ethanol precipitation at −20°C.

### Nucleosome core particle reconstitution

NCP reconstitutions were performed largely as previously described (16, 51). Briefly, histones H2A, H2B, H3, and H4 are individually resuspended at 2 mg/mL in Unfolding Buffer (6 M guanidine-HCl, 20 mM Tris pH 7.5, 5 mM DTT). Most histone complexes were refolded as an H2A/H2B dimer and H3/H4 tetramer, except for complexes with K4/9/14/18/23/27R-H3 and tailless-H3, which were refolded as an H2A/H2B/H3/H4 octamer. All complexes were refolded at 1 mg/mL histone via dialysis into a High Salt Buffer (2 M KCl, 20 mM Tris pH 7.5, 1 mM EDTA, 0.5 mM benzamidine, 1 mM DTT). Each histone complex was purified via FPLC using a HiLoad 16/600 Superdex 200 column (GE Healthcare), and purity was assessed using 18% SDS-PAGE gel electrophoresis. Prior to the final NCP reconstitution, purified 147-bp Widom 601 DNA was resuspended in High Salt Buffer at a final concentration of 8-11 μM. The final NCP reconstitution was performed at a molar ratio of 1:1:2.2 or 1:1:2 in terms of 147-bp Widom 601 DNA:H3/H4 tetramer:H2A/H2B dimer. For samples with K4/9/14/18/23/27R-H3 and tailless-H3, the final NCP reconstitution was performed at 1:1:0.2 molar ratio of 147-bp Widom 601 DNA:H2A/H2B/H3/H4 octamer:H2A/H2B dimer. The NCP reconstitution was performed via slow desalting with an exponential gradient followed by overnight dialysis into No Salt Buffer (20 mM Tris pH 7.5, 1 mM EDTA, 0.5 mM benzamidine, 1 mM DTT). NCPs were purified via a 10-40% sucrose gradient (BioComp Instruments). NCP purity was assessed via sucrose gradient profile (absorbance at 260 nm) and 5% native-PAGE. Additionally, histone integrity was assessed using 18% SDS-PAGE prior to use in experiments.

### Expression and purification of H3 tail peptides

Unmodified H3(1-44) was used as an extended histone tail peptide sequence. The following H3(1-44) mutants were also used: R2/8Q, R17/26Q, R2/8/17/26Q, K4/9Q, K14/18/23/27Q, K4/9/18/27Q, K4/9/14/18/23/27Q, R2/8/17/26K, and K4/9/14/18/23/27R. Mutant H3 tail peptides were generated using Q5 Site-Directed Mutagenesis Kit (New England Biolabs). Peptides were expressed and purified as previously described (16). Briefly, each peptide was expressed from a pET3a vector in BL21(DE3) competent *E. coli* (New England Biolabs). Following cell lysis, the H3 tail peptides in the soluble portion were purified using Q-Sepharose Fast Flow resin (GE Healthcare) and a Source 15S column (GE Healthcare) via FPLC (Bio-Rad). Following lyophilization, peptides were purified in S30 Buffer (50mM KPi pH 7, 50mM KCl, 0.5mM EDTA) via FPLC using a HiLoad 16/600 Superdex 30 pg column (GE Healthcare). Peptides were further purified using a PROTO 300 C18 column (Higgins Analytical) via HPLC (Shimadzu). ESI mass Spectrometry (Finnigan LTQ, Thermo Electron Corporation, San Jose, CA) was used to confirm intact masses for H3 tail peptides. Peptides were lyophilized and stored at −20°C prior to experiments.

### Preparation of plates for phase separation assays

Prior to phase separation assays, each multiwell plate was washed with 1% Hellmanex at 30°C for 40 min, washed extensively with autoclaved ddH_2_O, and left to air dry. Each plate was then treated with Sigmacote (Sigma-Aldrich) for 10 min, washed extensively with water, and left to air dry.

### NCP stock preparation for phase separation assays

NCP stocks were prepared in MOPS Buffer [20 mM MOPS pH 7 (8 mM NaOH to pH), 0.1 mM EDTA] using a 10 kDa MWCO centrifugal filtration unit (Millipore Sigma) and were stored at 4°C for a minimum of 12 hours before checking stock concentration. To determine stock concentration, a secondary NCP stock was made into MOPS Buffer. NCP concentration was determined spectrophotometrically from a 1:100 dilution of the secondary stock into 2 M KCl using a NanoDrop (Thermo Fisher). Concentration was calculated using the 147-bp Widom 601 DNA extinction coefficient at 260 nm (ε_260_ = 2,312,300.9 M^−1^cm^−1^). The primary stock concentration was then back-calculated.

### Turbidity assay

All components [NCP, H3(1-44), 601 DNA, MgCl_2_ stock, and KCl stock] were prepared in MOPS buffer [20 mM MOPS pH 7 (with 8 mM NaOH to pH), 0.1 mM EDTA] and warmed to room temperature before conducting each assay. Final buffer conditions were 20 mM MOPS pH 7 (8 mM NaOH to pH), 0.1 mM EDTA, 150 mM KCl, and 2 mM MgCl_2_. Note that the addition of 2 mM MgCl_2_ is distinct from our previously reported phase separation assays (16). Most assays were run in µClear 96-Well Half Area Microplates (Greiner Bio-One) sealed with Optical Adhesive Film (Thermo Fisher Scientific). Absorbance measurements were recorded using a FlexStation 3 plate reader (Molecular Devices) at 600 nm (A_600_) for 60 minutes at one-minute intervals. Data were recorded with a set temperature of 25 °C, which had an actual experimental range of 24.9-25.6 °C (average with standard deviation of 25.3 ± 0.1 °C for NCP samples). The maximum A_600_ value measured within the first 10 minutes of the time course was used in subsequent analyses. Data were plotted using Igor Pro (Wavemetrics) and R Statistical Software (v4.2.2; R Core Team 2022).

For assays with NCP, components were mixed in the order of MOPS buffer, KCl, MgCl_2_, and with the final addition of concentrated NCP stock to initiate the reaction. Experiments were conducted at NCP concentrations of 0, 10, 25, 50, 100, 200, and 300 µM. A total of three replicates were collected for each sample, with varied NCP concentration order between replicates. Assays were conducted at a final volume of 36.5 μL in µClear 96-Well Half Area Microplates (Greiner Bio-One) or at a final volume of 75 μL (for matched sample height) in µClear 96-Well Full Area Microplates (Greiner Bio-One).

For H3 tail and DNA, components were mixed in the order of MOPS buffer, KCl, MgCl_2_, 147 bp 601 DNA, and with the final addition of concentrated H3(1-44) stock to initiate the reaction. Assays were performed at 50 μM H3 tail peptide with increasing DNA concentrations (0, 0.625, 1.25, 2.5, 5, 10, 12.5, 15, 25 μM) and a final volume of 20 μL in each well.

### Brightfield microscopy

Images were acquired using Cytation10 Confocal Imaging Reader (Agilent) in brightfield mode with a 60x objective, collecting from −30 µm to 200 µm in 2 µm steps. Samples were prepared in a similar manner to those for turbidity assays. All components [NCP, H3(1-44), 601 DNA, MgCl_2_ stock, and KCl stock] were prepared in MOPS buffer [20 mM MOPS pH 7 (with 8 mM NaOH to pH), 0.1 mM EDTA] and warmed to room temperature before conducting each assay. Final assay buffer conditions were 20 mM MOPS pH 7 (8 mM NaOH to pH), 0.1 mM EDTA, 150 mM KCl, and 2 mM MgCl_2_. Assays used µClear 96-Well Half Area Microplates (Greiner Bio-One) sealed with Optical Adhesive Film (Thermo Fisher Scientific). Final experimental well volumes were 20 or 36.5 μL. The microscopy images were collected at room temperature with an experimental range of 27.4-28.2 °C for peptide-DNA assays and 27.4-29.0 °C (average with standard deviation of 28.0 ± 0.3 °C) for NCP assays.

The NCP was pipetted and imaged in two ways in order to capture faster-settling condensates. The first method pipetted concentrations 0 to 300 µM in order and imaged from 0 to 300 µM. The second method pipetted concentrations 0 to 300uM in order and imaged from 300 to 0 µM. NCP microscopy replicates were conducted with n = 4 for unmodified and n = 3 for all mutants, which includes two biological replicates for all constructs except the two single mutants (R2Q and R8Q). Additional concentrations were filled in at 5 or 20 µL volumes in µClear 384-well small volume LoBase or 96-Well Half Area Microplates (Greiner Bio-One), collected in triplicate with a single biological replicate. At 300 µM NCP, phase-separated samples of R2/8/17/26Q-H3-NCP separated into two distinct layers by the time automated data collection began, with the lack of visible droplets falsely suggesting a lack of phase separation. Thus, for this mutant at 300 µM, an additional single-well experiment was collected to shorten the experiment dead time as much as possible. This single-well imaging is used in **Figure 1D**.

**Figure 1.**
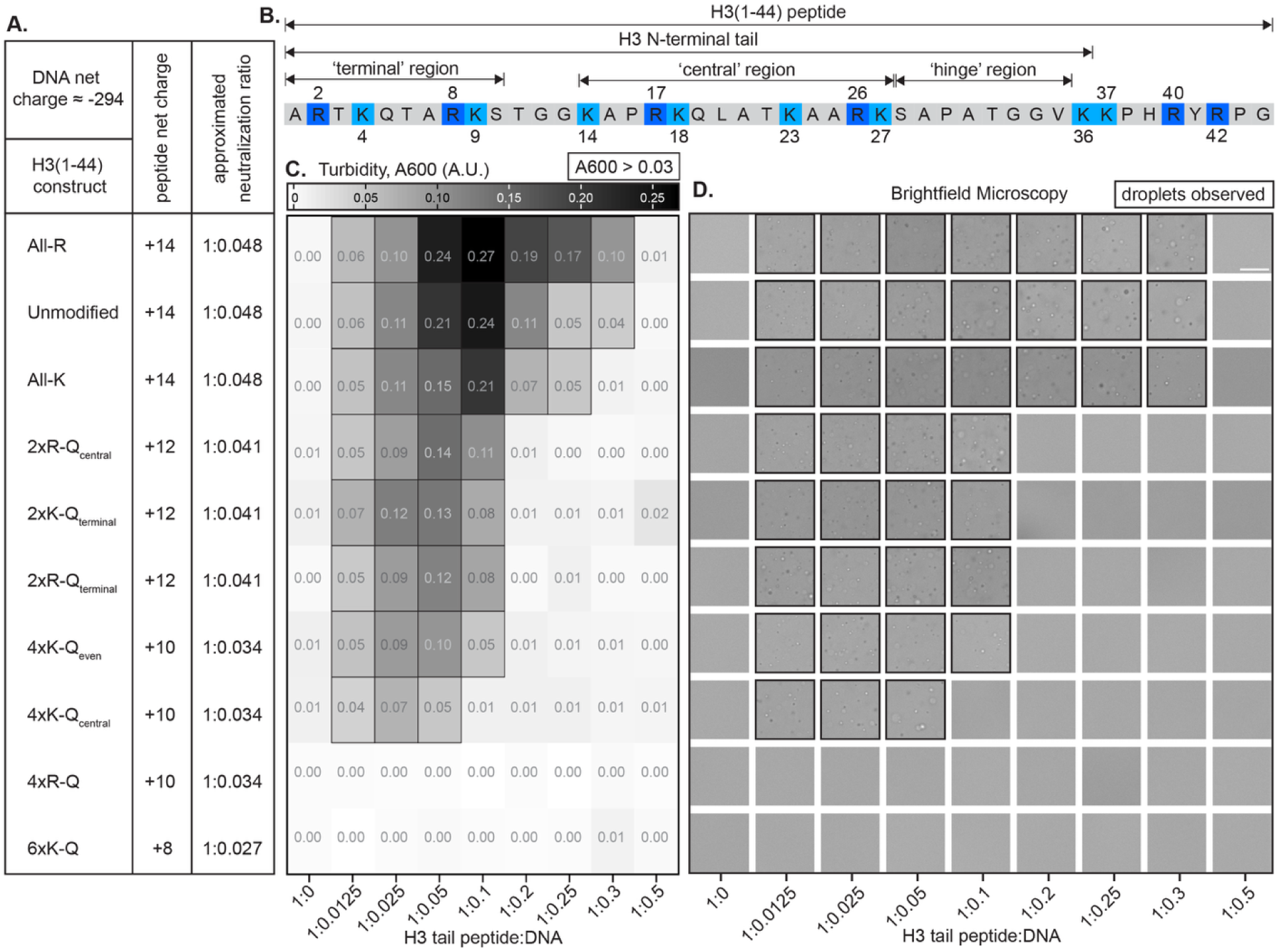
Phase separation of H3 tail peptide with DNA is modulated by perturbation of basic residues. **A.** Mutants of H3(1-44) are summarized and shown explicitly in **Supplementary Figure 1**. The net charge and estimated charge-neutral molar ratio of peptide:DNA are tabulated for each mutant. **B.** Sequence of H3 peptide, H3(1-44), which includes the H3 tail. Arginine and lysine residues are highlighted in shades of dark and light blue, respectively. **C.** Turbidity (absorbance at 600 nm) values (average of n=3) are shown in greyscale for unmodified and mutant versions of H3(1-44) at 50 µM with increasing amounts of 147-bp Widom 601 DNA, shown as molar ratio. Turbidity is plotted as a heatmap ranging from 0.00 (white) to 0.27 (black). Values greater than 0.03 are outlined with a black square to indicate light scattering due to droplet formation. Data were recorded with a set temperature of 25 °C, which had an actual experimental range of 24.9-25.6 °C. **D.** Brightfield microscopy images were collected at the same molar ratios as in (**C**). The scale bar is 25 µm. Images are outlined in black to indicate droplets. See **Supplementary Figure 2** for an alternative representation of turbidity data and larger images. Microscopy images were collected at room temperature with an experimental range of 27.4-28.2 °C. Construct identity in (**C**) and (**D**) aligns with the table in (**A**).

Images for figures were made using Gen5 (Agilent). Images were further analyzed using ImageJ version 2.1.0/1.54f. In an attempt to objectively define the presence of droplets in brightfield images, a pseudo-coefficient of variation (CV) was calculated. The total pixel intensity values of a standard deviation z-projection and an average intensity z-projection (from 0 to 200 µm) were determined for each NCP concentration. The reported CV was taken as the ratio of the intensities of the standard deviation projection and the average intensity projection, multiplied by 100. The CV values were averaged across replicates and plotted (**Supplementary Figure 4A**) using R Statistical Software (v4.2.2; R Core Team 2022). A CV cutoff of 5.0 was used to delineate phase separation. These values agree with a visual assessment of droplets. For H3 tail peptide and DNA, only one set of microscopy images was obtained, which were visually assessed for droplets.

### Fluorescence microscopy assay

For fluorescence imaging of NCP samples, YOYO-1 dye (Invitrogen) was added to NCP stocks at a 1:100 molar ratio with a minimum incubation time of 16 hours prior to inducing phase separation and imaging. Samples were prepared at 200 μM NCP in a final volume of 4 μL. Final buffer conditions were 20 mM MOPS pH 7 (8 mM NaOH to pH), 0.1 mM EDTA, 150 mM KCl, and 2 mM MgCl_2_. Samples were imaged at room temperature in µClear 384-well small volume LoBase microplates (Greiner Bio-One) sealed with optical adhesive film (Thermo Fisher Scientific). Microscopy was performed on a Nikon Eclipse Ti2-E inverted spinning disc microscope equipped with a Yokogawa confocal scanner unit (CSU-W1), 4 laser lines (405/488/568/633), a Hamamatsu ORCA-Flash4.0 V3 sCMOS Camera system with an 82% quantum efficiency chip and a water immersion dispenser. The system is run using the NIS-Elements Advanced Research package software (version X). DIC and confocal images were collected with a z-step size of 2 µm using a 60x water objective (SR Plan Apo IR). The field of view for the camera is 2048 x 2044 pixels with resolution 0.16 µm/pixel for the 60x water objective. Images were processed using Fiji and ImageJ version 2.1.0/1.54f.

### NCP sample recovery, quality assessment, and re-use

Following assays with non-fluorescently labeled NCPs, samples were pooled and diluted to reduce salt and dissipate NCP phase separation. Both pre-assay and post-assay NCP samples were run on 5% native-PAGE and 18% SDS-PAGE. Samples were then concentrated and exchanged into MOPS buffer using a 10 kDa MWCO centrifugal filtration unit (Millipore Sigma). Following gel confirmation and exchange, the samples were reused in replicate phase separation assays as done previously (16).

### Video particle tracking (VPT) microrheology of NCP samples

VPT measurements were performed on phase-separated samples of NCP variants prepared at a final NCP concentration of 200 μM in 20 mM MOPS pH 7 (8 mM NaOH to pH), 0.1 mM EDTA, 150 mM KCl, and 2 mM MgCl_2_. Yellow-green carboxylate-modified polystyrene beads of 1 µm size were mixed during sample preparation. These beads were confirmed to partition into the dense phases of phase-separated NCP samples (**Supplementary Video 1**) and were used as probe particles for VPT measurements. The samples of volume 5 or 10 μL were sandwiched between a glass slide and coverslip treated with 5% w/v PEG in 5% ethanol. The samples were surrounded by mineral oil to prevent evaporation. The prepared sample sandwich was placed on a temperature-controlled stage (Instec) attached to a Zeiss Primovert inverted microscope. Temperature was controlled using the thermal stage, while the actual temperature was determined using a thermocouple. The samples were allowed to fuse before imaging and were rested on the microscope for 30-60 minutes before data collection. The bead motion inside the NCP dense phases was captured at 24 °C in the form of videos using a 40x air objective lens and Teledyne FLIR Blackfly S USB3 CMOS camera for 1000 frames with 100 ms exposure time. The acquired videos were processed using Fiji (version 1.54f) (52). The probe particles were tracked to get their trajectories using the Trackmate (53) software plugin in Fiji. During tracking, intensity and quality filters were used to avoid tracking aggregated particles.

The analysis routine for estimating the mean square displacement (MSDs) from particle trajectories is detailed in our previous work (54, 55). In brief, the center of the mass trajectory of the bead motion was utilized to correct the bead trajectories for any possible drift during video acquisition. The center of the mass trajectory was estimated from the velocities of the bead using

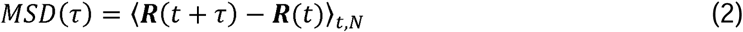

where ***X_COM_*** is the center of mass vector calculated for *k*_th_ frame, *j* is the individual frame, and *N_j_* is the number of particles in the frame *j*. ***X*_0_** is the center of the mass vector of the first frame. After correction, the obtained position vector of the particle ***R*** can be used to estimate the MSDs using

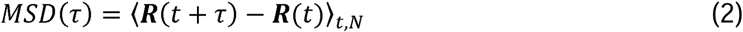

where *τ* is the lag time. We used our custom Python scripts, also deposited and can be found on GitHub (https://github.com/BanerjeeLab-repertoire/Material-properties), to compute MSD across various lag times *τ*. The ensemble average MSD was then determined based on data from approximately 20–50 individual beads. The MSD was then fitted (54, 55) using

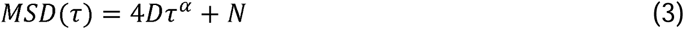

to obtain the diffusion coefficient *D* of the beads. Here, *α* is the diffusivity parameter which signifies the nature of bead diffusion inside the dense phases of NCP samples. The ensemble-average MSDs for which the fitting yielded *α* =1 were used to estimate the viscosity *η* using the Stokes-Einstein equation (54, 55)

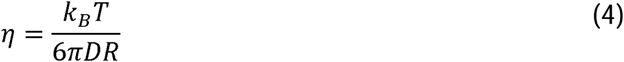

where *K_B_* is the Boltzmann constant, *R* is the radius of the particles, *D* is the diffusion coefficient of beads obtained by fitting the ensemble-average MSDs to Eq. (3) and *T* is the temperature of the sample.

The videos were recorded at 9-10 randomly chosen positions of the dense phases of NCP samples, and data were collected on three technical replicates for each NCP mutant. Statistical analyses were performed using R Statistical Software (v4.2.2; R Core Team 2022). To determine variance within viscosity measurements, a one-way ANOVA was initially used. Measurements obtained were determined to be significant with a p-value of 1.11E-64 (n=3). To determine which NCP viscosity measurements are significant to one another, a post-hoc Tukey’s test was used where a p-value of < 0.05 was deemed significant for the pairwise comparisons between the NCPs. Data were plotted using R.

### NMR sample preparation

Purified NCPs reconstituted with ^15^N-H3 (Unmodified, 4xR-Q, All-K, or All-R) were exchanged into MOPS buffer [20 mM MOPS pH 7 (with 8 mM NaOH to pH), 0.1 mM EDTA] using consecutive rounds of concentration and dilution in 10k MWCO centrifugal filtration devices (Millipore Sigma). NCP stock concentration was determined spectrophotometrically via absorbance at 260 nm after diluting into 2 M KCl.

Phase separation was induced by diluting to 400 µM NCP in final solution conditions of 18 mM MOPS pH 7 (with 7.2 mM NaOH to pH), 0.09 mM EDTA, 150 mM KCl, 2 mM MgCl_2_, and 10% D_2_O. The sample was transferred to a 1.7 mm NMR tube (Bruker) and left to separate into two layers overnight. The top (light phase) layer was then transferred to a second 1.7 mm NMR tube. Care was taken to ensure the NMR coil volume was filled with either dense or light phase.

### NMR data collection

Data were collected on a Bruker Avance III HD 600 MHz spectrometer with a 1.7-mm cryogenic probe at 298 K and running Topspin 3.5pl7. For chemical shift analysis, ^1^H-^15^N HSQC spectra were collected with 16 scans, 1024 (^1^H) x 300 (^15^N) total points, acquisition times of 51 ms (^1^H) and 75 ms (^15^N), and spectra widths of ~17 ppm (^1^H) and ~33 ppm (^15^N). All nuclear spin relaxation experiments were collected with 1600 (^1^H) x 228 (^15^N) total points, acquisition times of 102 ms (^1^H) and 84 ms (^15^N), and spectra widths of 13 ppm (^1^H) and 22 ppm (^15^N). {^1^H}-^15^N steady-state heteronuclear nuclear Overhauser effect (hnNOE) data were collected using the Bruker pulse sequence hsqcnoef3gpsi with the reference and saturated spectra interleaved. Experiments were collected with 32 (for dense phase samples), 72 (for 4xR-Q-H3 light phase sample), 88 (for All-R light phase), or 96 (for unmodified-H3 and All-K-H3 light phase samples) scans and an interscan delay of 5 s. The dense phase hnNOE experiments were collected at the start of the experimental queue and again 13-18 days later to check for sample aging. The data were unchanged within experimental error, and data from the initial hnNOE experiment were used in figures. Longitudinal (R1) ^15^N relaxation experiments were collected with total relaxation loop lengths of 0.033 s, 0.195 s, 0.390 s, 0.585 (x2), 0.813 s, 1.301 s, and 1.951 s in randomized order. Experiments were collected with 24 (for dense phase samples), 32 (for R2/8/17/26Q-H3 light phase sample), or 40 (for unmodified-H3 and R2/8/17/26K-H3 light phase samples) scans and an interscan delay of 2 s. Transverse (R2) ^15^N relaxation experiments were collected with an effective CPMG field of 500 Hz and total relaxation CPMG loop lengths of 0.0044 s, 0.013 s, 0.026 s, 0.044 s (x2), 0.062 s, 0.084 s, and 0.110 s. Experiments included temperature compensation blocks and were collected in randomized order, with the same number of scans as for R1, and with an interscan delay of 2.5 s. ^1^H-^15^N HSQC spectra were collected between data series to confirm sample integrity. For ^15^N-All-R-H3-NCP, 1D versions of the R1 and R2 experiments were collected with 256 scans at the same delay times as for the 2D versions.

### NMR data analysis

NMRPipe and CcpNmr Analysis 2.5.2 were used within NMRbox for processing and analysis, respectively, of NMR data (56–58). Data were plotted using Igor Pro (Wavemetrics). Amide assignments for NCPs with unmodified H3 are from BMRB entry 50806. These assignments were transferred to H3 mutants via inspection and comparison to previously published spectra (59). Chemical shift differences (Δ*δ*) between solution phases were calculated from the differences in amide proton (Δ*δ_H_*) and amide nitrogen (Δ*δ_N_*) chemical shifts using: 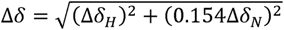 (60). Spectra were processed using a cosine-squared window function for both dimensions, and each dimension was doubled in size by zero-filling twice and rounding to the nearest power of two. HnNOE values for each residue were calculated as the peak height ratio between the saturated and reference spectra, and errors were calculated via standard error propagation of the spectral noise as determined by CcpNmr Analysis 2.5.2. R1 and R2 relaxation rates were determined by fitting peak heights from the relaxation series to a single-exponential decay without offset. Errors were determined from the covariance matrix within CcpNmr Analysis 2.5.2. As discussed for previous analyses, there is the caveat resulting from the low chemical shift dispersion of IDRs that relaxation rates can be convolution by nearby peaks in dense spectral regions (59). For the ^1^H-1D versions of R1 and R2 experiments, the amide backbone region was integrated in CcpNMR Analysis 3.3.2.1 and fit using Igor Pro (Wavemetrics) to a single-exponential decay without offset where the reported error is the estimated standard deviation of the fit coefficient.

### Coarse-grained molecular simulations of H3 tail with DNA fragments

We employed a one-bead hydrophobicity scale (HPS) model to calculate the phase diagram and viscosity of H3 tail-DNA co-condensates. The H3 tail sequence is the same as the peptide tested experimentally (ARTKQTARKSTGGKAPRKQLATKAARKSAPATGGVKKPHRYRPG). A 20-thymine (T) strand complementary base-paired with adenine (A) was used as the DNA sequence. The two-component system comprises double-stranded DNA and H3 peptides with 40 and 44 beads, respectively. The beads in the DNA chain represent adenine-base (A) and thymine-base (T) nucleotides, and the beads in the peptide represent amino acids in H3(1-44). The non-bonded interactions between residues *i*, *j* are modeled via the Ashbaugh-Hatch potential, and salt-screened electrostatic interactions are modeled via the Debye-Hückel potential:

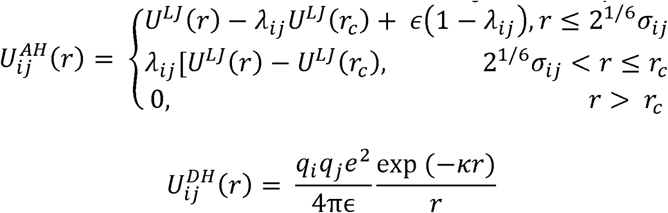

where *ɛ* = 0.8368 kJ/mol, r_c_= 4 nm, and U^LJ^ is the Lennard-Jones potential 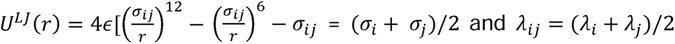 and *λ_ij_* = (*λ_i_* + *λ_j_*)/2 are arithmetic averages of monomer size and hydrophobicity value, respectively. The pair potential with *λ* = 0 consists of only the repulsive term equivalent to the Weeks-Chandler-Andersen functional form. We use *σ* values of amino acids from van der Waals volumes and *λ* values of amino acids from the recently proposed CALVADOS 2 parameters (61). In the Debye-Hückel potential, *q* is the charge number of particles and 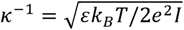 is the Debye length. In our simulations, the ionic strength of the electrolyte *I* = 0.17 M. The temperature-dependent dielectric constant used in our simulations has the following empirical relation: 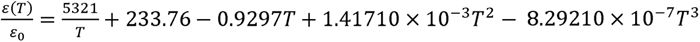. All coarse-grained beads are connected via bonded interactions modeled by harmonic potentials 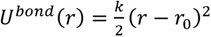 where k = 1000 kJ/mol/nm^2^, r_0_ = 0.5 nm for DNA, and r_0_ = 0.38 nm for peptide. In dsDNA, there are additional base pair interactions and stacking interactions between adjacent base-pair groups. The base pair interaction is represented by a repulsive Morse potential, *U_bp_* = *D_exp_*(−*α*(*r* − *l*_0_)) with *D* = 472 kJ/mol, *α* = 2.876 nm^−1^ and *l*_0_ = 1.243nm. The stacking interaction is modeled using a Lennard-Jones potential, characterized by *ɛ* = 49.2 kJ/mol, *σ* = 0.2938 nm, and a cut-off distance *r_C_* = 5.65 nm. We have applied OpenMM (v7.7) for H3 tail-DNA condensate simulations.

We performed coexistence simulations in a slab geometry at different *T* for characterizing their phase diagrams. The number of dsDNA was set at N_DNA_ = 180. The number of H3 tail peptides was set around N_H3_ = 520 − 720. These simulations were carried out at each *T* for a duration of 5 × 10^7^ – 7 × 10^7^ steps, and the dense phase and dilute phase concentrations were computed based on the last 2 × 10^7^ steps of the simulation trajectory. We estimated the critical temperatures by fitting the phase diagram using the law of coexistence densities and the critical concentrations by assuming the law of rectilinear diameter holds.

We ran condensate simulations in a cubic box for viscosity measurement. The number of dsDNA was set around N_DNA_ = 84-180. The number of H3 tail peptides was set around N_H3_ = 254 − 426 to keep the approximate charge neutrality of the co-condensates. The total number of the CG monomers was kept at about 2 × 10^4^ to eliminate the finite-size effect. The initial length of the cubic box was set at 40 nm, which is much larger than the equilibrium size. The box was then shrunk in the subsequent *NPT* simulations. After energy minimization, we used the *NPT* ensemble with a timestep of 0.01 ps, to equilibrate all systems using an additional 5 × 10^7^ steps. Finally, we fixed the box size as the average volume in *NPT* simulations and carried out additional *NVT*simulations for 1 × 10^7^ steps.

## RESULTS

### Design of H3 tail basic residue mutants to probe nucleosome phase separation

We set out to investigate the role of charge neutralization of H3 tail basic residues and the distribution of these neutralizations along the tail on nucleosome phase separation. Charge-modulation of the histone tails is expected to perturb the balance between multivalent intra- and inter-nucleosomal tail-DNA interactions (24) and, in turn, phase separation propensity and condensate properties. Furthermore, the most N-terminal basic residues in the tail are anticipated to have greater potential to form inter-nucleosomal interactions and facilitate phase separation than the basic residues in the center of the tail sequence.

Here, we focus on six lysine and four arginine residues within the H3 tail that can promote phase separation through inter-nucleosomal electrostatic interactions with DNA. To examine the position-dependent effects of perturbing these basic residues, we divided the H3 tail into three regions: the **‘terminal’ region**, comprising residues preceding the first TGG motif; the **‘hinge’ region**, corresponding to the uncharged segment that contains the second TGG motif and precedes the histone core; and the **‘central’ region**, spanning between the ‘terminal’ and ‘hinge’ regions (**Supplementary Fig. 1**). Neutralizing mutations of basic residues to glutamine in the terminal and central regions were introduced as PTM mimetics of arginine citrullination and lysine acetylation. We designed H3 mutants (**Supplementary Figure 1**) to separately neutralize arginines and lysines within the terminal region (R2/8Q, referred to as 2xR-Q_terminal_, and K4/9Q, referred to as 2xK-Q_terminal_) and central region (R17/26Q, referred to as 2xR-Q_central_, and K14/18/23/27Q, referred to as 4xK-Q_central_). Additionally, we designed H3 mutants with neutralization of the two terminal region arginines individually (R2Q and R8Q), all four arginine residues (R2/8/17/26Q, referred to as 4xR-Q), the four lysine residues adjacent to arginine residues (K4/9/18/27Q, referred to as 4xK-Q_even_), and all six lysine residues across the terminal and central regions (K4/9/14/18/23/27Q, referred to as 6xK-Q). To test the importance of H3 tail basic residue identity, we also designed two mutants to convert all basic residues across the terminal and central regions of the H3 tail to only one type: R2/8/17/26K-H3 (All-K-H3) and K4/9/14/18/23/27R-H3 (All-R-H3).

### Sequence modifications to the H3 tail modulate tail-DNA condensate formation

To probe the effect of H3 tail modifications, we started with a simplified two-component system containing purified H3 tail peptide [residues 1-44, H3(1-44)] and the 147-bp Widom 601 DNA sequence (**Figure 1A-B**). Previously, we demonstrated that the phase separation behavior of H3 tail peptide with DNA is similar to the reentrant condensation observed for other systems with multivalent interactions between nucleic acids and arginine- and lysine-rich intrinsically disordered peptides (16, 44, 45, 62–64). Using microscopy and turbidity measurements, we determined the phase boundary for this two-component system and compared it to the estimated charge neutralization ratio for H3(1-44) mutants and 147-bp Widom DNA, which is predicted to display the maximum turbidity if the system follows complex coacervation (**Figure 1A, 1C, 1D**). Overall, increasing the number of neutralizations narrows the condensate formation regime with a subtle shift in maximum turbidity to lower ratios of DNA to peptide (**Figure 1C-D, Supplementary Figure 2)**. Phase separation near the estimated charge neutralization ratio is seen for all H3 tail peptides except with 4xR-Q and 6xK-Q mutations, for which no phase separation is observed. In contrast to 4xR-Q-H3(1-44), the two 4xK-Q mutants phase separate with DNA, supporting a differential effect between modifications to arginine and lysine. The similar effect of 2xR-Q_terminal_ and 2xR-Q_central_ mutations suggests no clear position dependence within this simplified two-component system, although a mild narrowing of the phase boundary is observed between the two 4xK-Q mutants. Unmodified-, All-K-, and All-R-H3(1-44) have the same net charge and display similar phase separation, but All-R-H3(1-44) is distinct in showing a broader transition via turbidity (**Figure 1C-D**, **Supplementary Figure 2A**). These results suggest a differential impact between arginines and lysines in driving condensate formation via H3 tail-DNA interactions, as well as a role for neutralization modifications in regulation of these condensates.

### Nucleosome core particle phase boundary shifts with basic residue neutralization in a position-dependent manner

We next asked whether this series of H3 modifications influences phase separation within the context of the nucleosome, where the H3 tails are only two of ten tails and are tethered to the core in close proximity to DNA. In this situation, we predicted a positional dependence of basic residue neutralization. To probe the effect of charge-neutralizing mutations of H3 tail basic residues on NCP phase separation (**Figure 2A-B**), we started by determining the saturation concentration (c_sat_) of each mutant-H3-NCP using turbidity and brightfield microscopy. These assays were conducted near room temperature (25.3 ± 0.1 °C and 28.0 ± 0.3 °C, respectively) and at physiologically relevant salt concentrations of 150 mM KCl and 2 mM MgCl_2_, for a range of discrete NCP concentrations from 0 to 300 μM.

**Figure 2.**
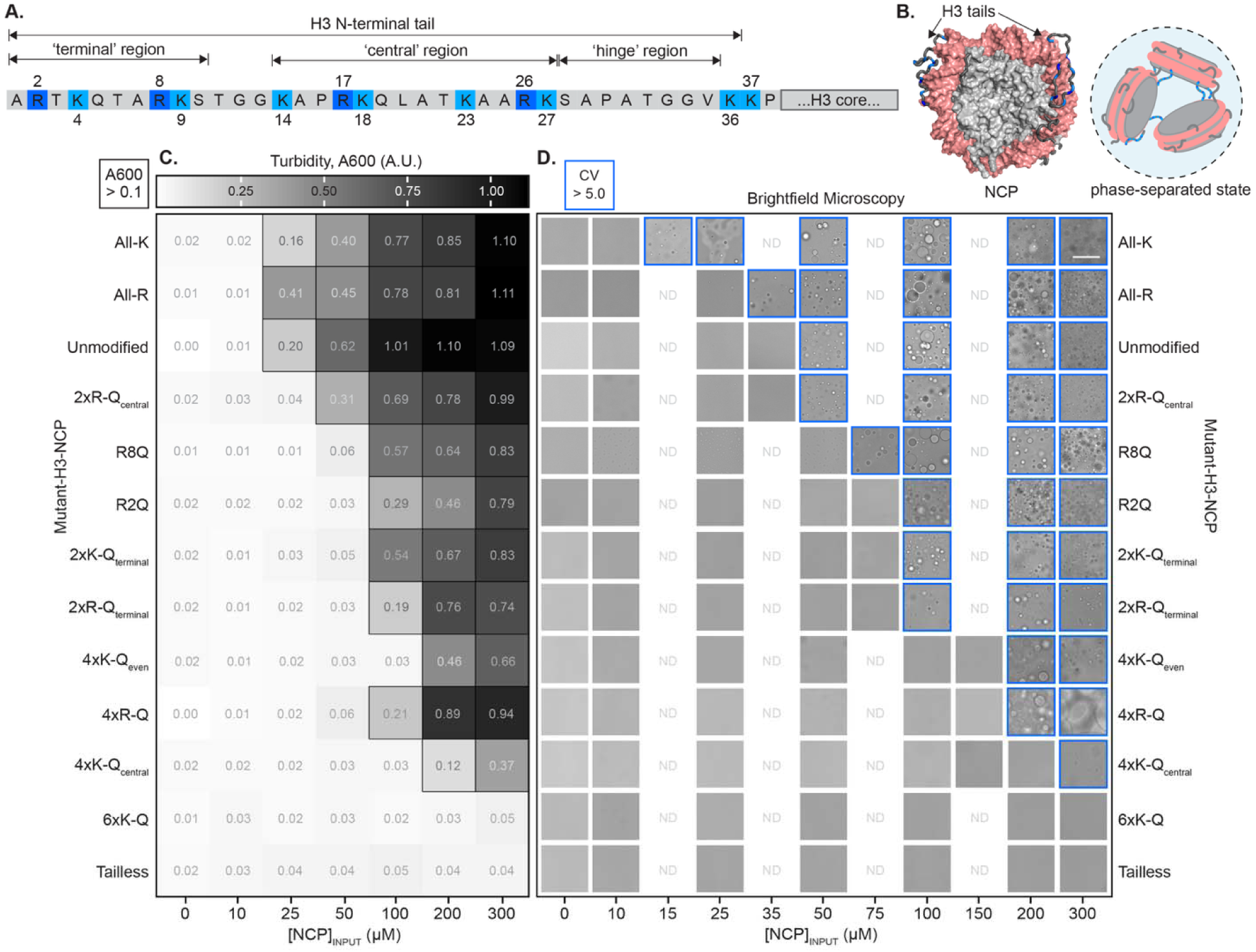
Nucleosome phase boundary shifts with perturbation of basic residues. **A.** Histone H3 tail amino acid sequence that protrudes from the nucleosome core. Arginine and lysine residues are highlighted in dark and light blue shades, respectively. **B.** NCP with basic residues of H3 tail colored according to (**A**). NCP conformation is from a molecular dynamics simulation end state (65). A cartoon illustrates that, while inter-nucleosomal tail-DNA interactions are a main driver of nucleosome phase separation, both intra- and inter-nucleosomal interactions are presumed to exist. **C.** Turbidity (absorbance at 600 nm) values (average of n=3) are shown in greyscale for NCP with unmodified or mutant H3, as labeled, and as a function of NCP concentration. Turbidity is plotted as a heatmap ranging from 0.01 (white) to 1.1 (black). Values greater than 0.1 are outlined with a black square to indicate light scattering due to droplet formation. Data were recorded with a set temperature of 25 °C, which had an actual experimental range of 24.9-25.6 °C. **D.** Brightfield microscopy images were collected for the same NCP concentrations as in (**C**), along with additional concentrations to improve c_sat_ determination. The scale bar is 50 µm. A coefficient of variation value (see Methods) was determined for each image. Values greater than 5.0 are considered significant and are outlined in blue to indicate droplets. Full field-of-view images are shown in **Supplementary Figure 4**. Microscopy images were collected at room temperature with an experimental range of 27.4-29.0 °C.

Unmodified-NCP was used as the point of comparison for all mutants. Droplets were observed via brightfield microscopy at unmodified-NCP concentrations of 50 μM and greater, which is also in agreement with sample turbidity (**Figure 2C-D**, **Supplementary Figures 3**-**4**, **Supplementary Tables 1-2**). Note that this boundary is shifted to lower concentrations from our previous publication due to the inclusion of MgCl_2_ (16). In agreement with previous results acquired in the absence of MgCl_2_ (16), joint removal of the terminal and central regions of the H3 tail (referred to as tailless-H3-NCP) abrogates phase separation at concentrations up to at least 300 µM (**Figure 2C-D**). We interpret these results to support the importance of the H3 tail in NCP phase separation under the experimental conditions tested.

A comparison of arginine and lysine mutants provides insight into the influence of charge neutralization of H3 tail basic residues on NCP phase separation as a function of identity, number, and position (**Figure 2C-D**, **Supplementary Figures 3-4**). When comparing arginine neutralizations, a dependence on number and position is observed: 2xR-Q_central_-H3-NCP has similar c_sat_ (50 µM NCP) as compared to unmodified-NCP while 2xR-Q_terminal_ and 4xR-Q shift the c_sat_ to higher concentrations (100 and 200 μM NCP, respectively). To further break down the effect of ‘terminal’ region arginines, we individually neutralized R2 and R8. R2Q produces a more pronounced shift in c_sat_ than R8Q (100 and 75 µM NCP, respectively) and is similar to the effect of the double neutralization. Lysine neutralizations exhibit a dependence on number: 2xK-Q_terminal_ shifts c_sat_ to 100 μM NCP while 4xK-Q_central_ and 4xK-Q_even_ shift c_sat_ to 200-300 μM NCP. The effects on c_sat_ are similar whether neutralizing two arginines or two lysines in the ‘terminal’ region; the effects of ‘central’ region neutralizations are confounded by the difference in the number of arginines and lysines (i.e., 2 vs. 4) in the ‘central’ region. When all six H3 tail lysines are neutralized (6xK-Q), we do not observe phase separation at the concentrations tested. A comparison of the conversion of all basic residues across the terminal and central regions of the H3 tail to only one type suggests that c_sat_ decreases to 15 and 35 μM for All-K and All-R, respectively (**Figure 2C-D**, **Supplementary Figures 3**-**4**), the only mutants tested to decrease c_sat_.

Together, these data suggest that the type (i.e., arginine vs. lysine), number (i.e., two vs four vs. six), and distribution (i.e., terminal vs. central) of basic residues neutralized play a role in defining nucleosome phase separation propensity. Results obtained within the NCP context exhibit trends similar to those observed in the simplified peptide–DNA system, but with an added dependence on residue position and a reduced sensitivity to residue number. It is somewhat surprising that the peptide–DNA model captures the overall trends of the more complex NCP system so well, given the absence of the other histone tails. Differential contributions of arginine and lysine to NCP phase separation propensity support that a mixture of arginine and lysine residues may be important for reasons beyond their distinct modifications and recognition by reader proteins.

### Basic residues within the histone H3 tail govern nucleosome condensate material properties

Next, we sought to understand how the neutralization of basic residues within the H3 tail influences the microenvironment of nucleosome condensates. Using temperature-controlled video particle tracking (VPT) microrheology (**Figure 3A**), we determined the viscosity of NCP condensates. Overall, neutralization of basic residues decreases the viscosity of NCP condensates, and the degree of neutralization correlates with the extent of enhanced condensate fluidity (**Figure 3B-C**, **Supplementary Tables 3-4**). We observe a decrease in viscosity when we increase the number of lysine neutralizations (compare unmodified at 110 ± 10 mPa s to 2xK-Q_terminal_- and 4xK-Q_even_-H3-NCP at 32 ± 8 and 13 ± 1 mPa s, respectively) and arginine neutralizations (compare unmodified to R2Q, R8Q, 2xR-Q_terminal_-, 2xR-Q_central_-, and 4xR-Q-H3-NCP at 39 ± 2 mPa s, 51 ± 4 mPa s, 26 ± 2 mPa s, 43 ± 5 mPa s, and 2.4 ± 0.5 mPa s, respectively). Neutralization of all four arginine residues within the H3 tail (4xR-Q-H3-NCP) leads to the lowest measured viscosity among the samples tested. Similar to c_sat_ (**Figure 2D**), there appears to be a dependence on position. Specifically, a more pronounced decrease in viscosity is observed for R2Q as compared to R8Q and with neutralization of two arginines in the ‘terminal’ region as compared to the ‘central’ region (compare 2xR-Q_terminal_- and 2xR-Q_central_-H3-NCP). Furthermore, we observe that the type of residue neutralized influences NCP condensate viscosity in some samples (compare 4xR-Q- and 4xK-Q_even_-H3-NCP). Interestingly, All-K-H3-NCP has decreased condensate viscosity (84 ± 4 mPa s) as compared to unmodified-NCP while All-R-H3-NCP is the one modification tested that increases the condensate viscosity (250 ± 30 mPa s). Collectively, the data reveal that modifications to basic residues enable tuning of NCP condensate viscosity across nearly two orders of magnitude. Notably, a global linear correlation exists between viscosity and c_sat_ on a natural logarithm scale with two exceptions: All-K- and 4xR-Q-H3-NCP (**Figure 3D**). These differences may stem from a complex interplay between hydrodynamic effects and sequence-encoded thermodynamic interactions in these systems (66).

**Figure 3.**
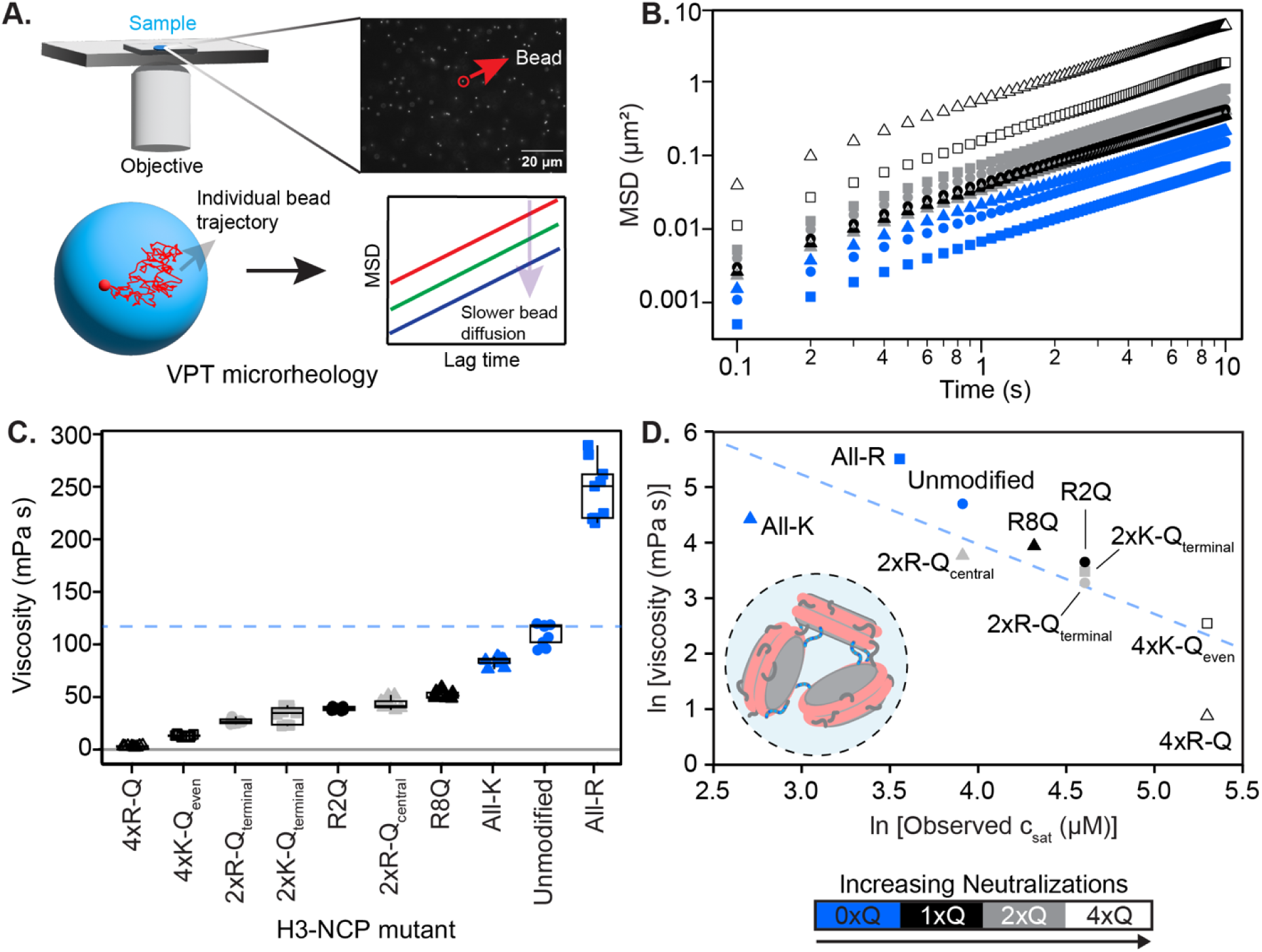
Nucleosome condensate viscosity is tuned by H3 tail charge patterns. **A.** Schematic representation of the video particle tracking (VPT) microrheology employed to determine the viscosity of condensates formed by unmodified- and mutant-H3-NCPs. The motion of 1-µm yellow-green fluorescent carboxylate-modified beads embedded inside condensates was tracked. Individual trajectories of the tracked beads were analyzed to estimate the mean squared displacement (MSD) of the beads. **B.** Ensemble-average MSD profiles of the tracked beads in condensates of unmodified- and mutant-H3-NCPs at 24 °C. A shift towards lower MSDs depicts slower diffusivity of the beads inside the dense phases of the NCP mutants. **C.** Viscosities of the condensates formed by unmodified- and mutant-H3-NCPs were estimated from the fit of the MSDs to the generalized Stokes–Einstein relation in the diffusive regime, using the power-law scaling of the form *MSD* = 4*Dτ^α^* (Methods section). In the box plot, the black center horizontal line marks the median, the grey box marks the interquartile range (IQR) from the 25^th^ to 75^th^ percentile, and the grey vertical lines mark the minimum (25^th^ percentile – 1.5*IQR) and maximum (75^th^ percentile +1.5*IQR). All viscosities are significantly different from unmodified NCP condensates as determined via ANOVA with Tukey post-hoc analysis. Additional significance comparisons are reported in **Supplementary Table 4**. **D.** Inverse correlation between the natural log of the viscosity and c_sat_ (as determined from brightfield microscopy in Figure 2D) of unmodified- and mutant-H3-NCP. The dashed blue line is a guide to the eye for the inverse correlation. The figure contains data from measurements at 9-10 randomly chosen positions of the dense phases over three independently prepared sample repeats. Data symbols are consistent across the figure.

### Phase behavior and viscosity of H3 tail-DNA condensates from coarse-grained simulations

The thermodynamic and material properties of H3 tail-DNA condensates were investigated using coarse-grained simulations (**Figure 4A-B**). To conduct the simulations, each amino acid of the histone tail, the same H3(1-44) peptide sequence as used experimentally (**Figure 1B**), and each nucleotide of a 20-bp poly(dT:dA) double-stranded DNA were coarse-grained into single beads. For the tail peptide, we use an established single-bead model called HPS-CALVADOS (61) in which the effect of residue identity on intermolecular attraction is encoded through the λ-parameter that quantifies the hydropathy scale of residues. In this parametrization, arginine has a substantially higher λ than lysine, reflecting its experimentally established stronger capability to engage in hydrogen-bonding, cation–π, and electrostatic interactions in crowded or partially dehydrated environments. As a result, arginine-rich sequences exhibit markedly stronger effective cohesion than their lysine-rich counterparts.

**Figure 4:**
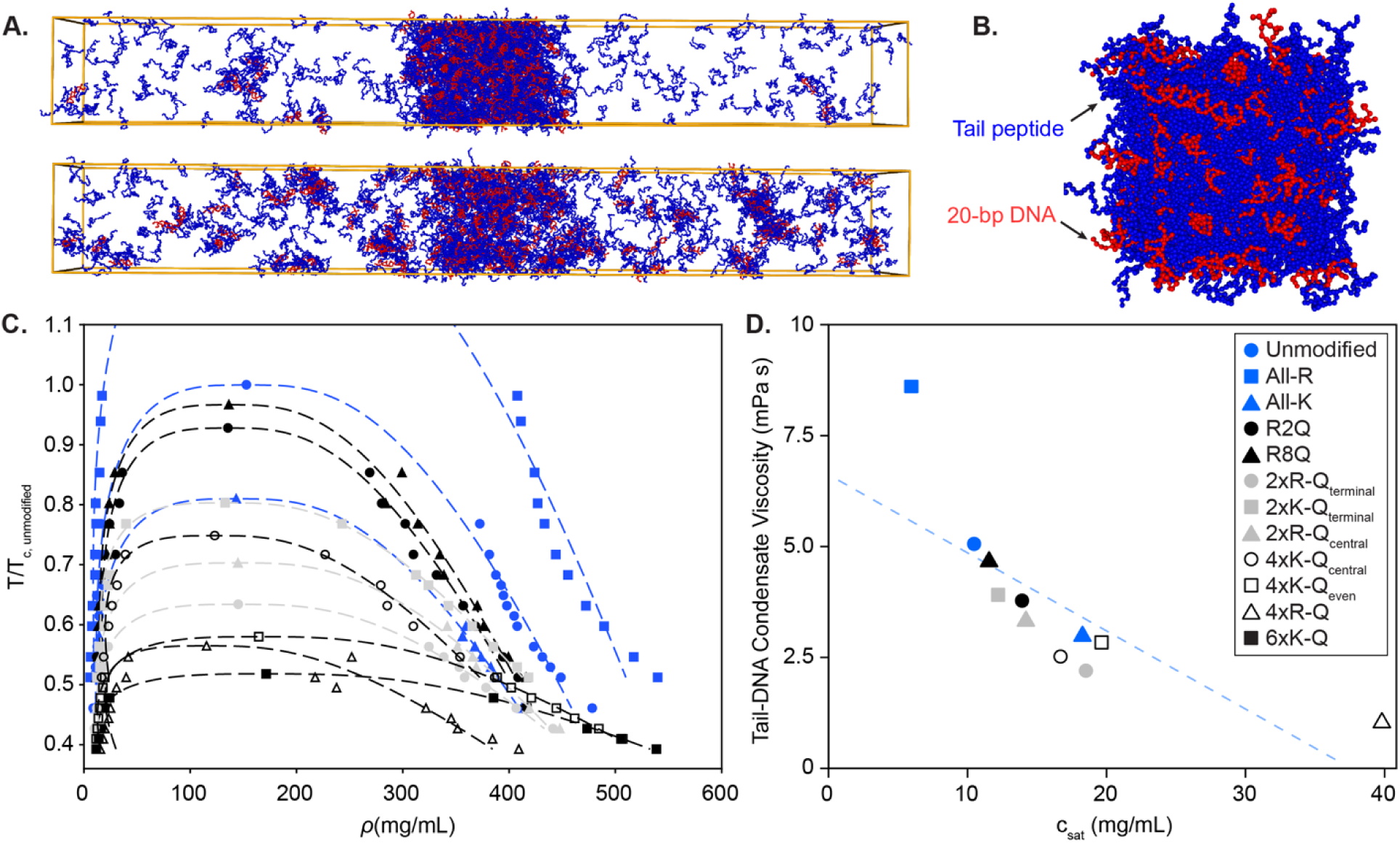
Phase behavior and viscosity of H3 tail-DNA condensates of unmodified and mutants from coarse-grained MD simulations. **A.** Snapshots of 4xR-Q-H3(1-44) (blue) with DNA (red) condensates at 250 K (upper) and 300 K (lower) from coarse-grained simulations, highlighting the transition between condensed and dilute phases. **B.** Snapshot from viscosity calculation simulations performed at 300 K, where the simulation box is systematically shrunk to achieve the condensed-phase density. **C.** Phase diagrams of unmodified- and mutant-H3 tail peptide with DNA, showcasing the critical temperature and concentrations associated with phase transitions. Temperature (T) is plotted as a ratio, relative to the critical temperature of unmodified-NCP (T_c, unmodified_). **D.** Calculated viscosity as a function of saturation concentration (c_sat_) derived from the phase diagram analysis of the unmodified- and mutant-H3 tail peptides. The dashed blue line is a guide to the eye for the inverse correlation. Symbol legend in (**D**) corresponds to (**C**) and (**D**).

The phase diagram (**Figure 4C**) shows broad agreement with the experimental results obtained with NCPs (**Figure 2C-D**). The All-R H3 tail mutant displays higher critical temperatures compared to the unmodified and all other mutants. With increasing arginine neutralization (compare unmodified, 2xR-Q_terminal_, and 4xR-Q), both the critical temperature and concentration decrease. The higher critical temperature of 2xR-Q_central_ compared to 2xR-Q_terminal_ suggests a positional effect, where the location of arginine neutralizations within the histone tail influences phase separation behavior and condensate stability. A comparison of the substitution of all arginines with either lysines (All-K) or glutamines (4xR-Q) shows a significant increase in the critical temperature for All-K, along with a broader phase separation boundary (i.e., the condensate concentration is higher at 300 K). The 4xR-Q mutant and its lysine corollary, 4xK-Q_even_, exhibit very similar values of critical temperatures as indicated by the fitting results. The 4xR-Q mutant exhibits the narrowest phase separation boundary compared to all other mutants and the unmodified peptide. The 6xK-Q mutant is identified by the coarse-grained model as having the lowest critical temperature, and under our experimental conditions, this mutant failed to form condensates either in the context of peptide-DNA systems or NCP (**Figure 1C-D**, **Figure 2C-D**).

Next, we quantified viscosities from coarse-grained simulations of the H3 tail–DNA system using the same sequence variants examined experimentally. In coarse-grained models, we have previously demonstrated that the microscopic origin of viscosity arises from the association and dissociation kinetics of sticky, positively charged residues (primarily arginine and lysine) at the DNA interface (55, 67). When these residues are clustered, their interactions with DNA become cooperative, increasing the effective dissociation energy. As a result, viscosity grows nonlinearly with the local density of high-affinity residues. The computational trends we obtain closely parallel the experimental measurements on NCPs (**Supplementary Figure 5**) and are fully consistent with this microscopic mechanism. All variants except All-R show reduced viscosity relative to unmodified H3, whereas All-R displays a dramatically elevated viscosity. That is because All-R represents a highly cohesive sequence: it contains nearly twice as many arginines as unmodified-H3(1-44), and the patterning of these residues is dramatically altered, creating several arginine repeats. No other constructs contain arginine dipeptides and at most one short window with two arginine residues in three consecutive positions. In contrast, All-R contains three arginine dipeptides and eight distinct 3-residue windows enriched in arginine. This dense, patterned clustering of strongly attractive residues produces a nonlinear amplification of cohesive interactions, leading to slower dissociation from DNA and therefore substantially higher viscosity. Within the remaining mutants, the 2xR-Q_terminal_ construct exhibits lower viscosity than the 2xR-Q_central_ mutant, and the 4xR-Q mutant shows the lowest viscosity of all. These results also support an inverse relationship between viscosity and c_sat_ across the studied constructs (**Figure 4D**, **Supplementary Table 5**), suggesting that both equilibrium stability and condensate dynamics are governed by the same underlying pattern of sequence-encoded interactions.

### H3 tails remain highly mobile in nucleosome condensates

As histone tail dynamics are proposed to be important in the histone language (59, 68, 69), we next investigated how these functional protein motions are perturbed by the condensate environment. While the condensate environment severely restricts local motions in some systems (70–72), several condensate systems have been reported to display high mobility of intrinsically disordered regions at the molecular level (73, 74). The measured viscosities of the NCP condensates (**Figure 3C**, **Supplementary Table 3**) are at the lower end of the values previously reported for biomolecular condensates (75), suggesting that the condensates are sufficiently fluid to permit experimental measurement of tail dynamics.

We used solution NMR spectroscopy to investigate the mobility of the H3 tails within NCP condensates. In these samples, only the H3 component of the NCP is ^15^N-labeled, and only the tail is visible by solution NMR with this labeling approach due to the large size of the complex (32, 76). Notably, backbone resonances were still observable for the nucleosomal H3 tails within the dense phase of a phase-separated sample, supporting that the tails retain a high level of mobility within condensates (**Supplementary Figure 6A**, left). However, resonances for residues towards the ends of the tail (see T3, K4, and V35) broaden almost beyond detection. Chemical shift differences (Δ*δ*) determined from ^1^H-^15^N HSQC spectra were minimal between the dense and light phases (**Supplementary Figure 6C**). This is commonly observed for IDPs within condensates (e.g., see (70, 71, 73)). The visibility of most H3 resonances in the dense phase spectra, along with the minimal changes in chemical environment, supports a model wherein the H3 tails remain disordered and bound to DNA (presumably via a mixture of intra-and inter-nucleosomal interactions) in a dynamic ensemble of conformations.

We employed NMR spin relaxation experiments to gain insight into nucleosomal H3 tail dynamics on the ps-ns timescale. On average, ^15^N-R1 values increased by 3% while ^15^N-R2 values increased by 71% in the dense phase as compared to the light phase (**Figure 5A-B**: open vs. closed circles, **Supplementary Figure 7**: grey vs. black). The hnNOE profiles for the two phases are the same within the experimental error (**Figure 5C**, **Supplementary Figure 8**), with the caveat that the signal quality is poor for ^15^N-unmodified-H3-NCP samples. Thus, while there are only minor, if any, differences between the two phases for ^15^N-R1 and hnNOE, the ^15^N-R2 values exhibit a strong dependence on phase (**Figure 5A-C**, **Supplementary Figures 7-8**). The R2 profile for residues 26-29 stands out as distinct between the two phases. The difference in R2 profile shape for R26, K27, S28, and A29 suggests the possibility of interesting conformational dynamics changes at the boundary between the ‘central’ and ‘hinge’ regions between the two phases, which we speculate could originate from differences between intra-and inter-nucleosomal interactions.

**Figure 5.**
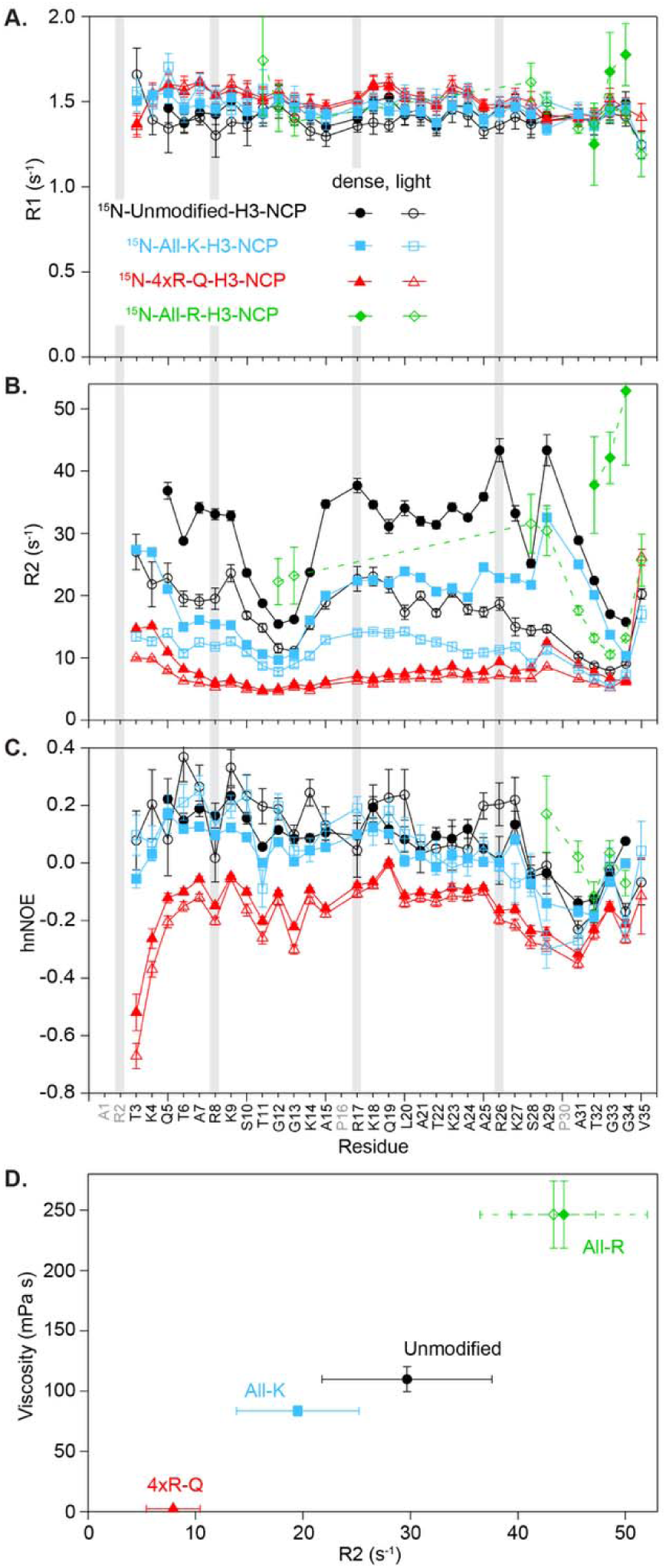
NMR fast-timescale dynamics indicate highly mobile H3 tails within nucleosome condensates. Plots are shown for ^15^N-R1 rates (**A**),^15^N-R2 rates (**B**), and hnNOE values (**C**) as a function of residue for unmodified- (black circles), All-K- (blue squares), 4xR-Q- (red triangles), and All-R- (green diamonds) H3-NCP. Data were collected on the light phase (open symbols) and dense phase (closed symbols) of NCP samples prepared in 20 mM MOPS pH 7, 150 mM KCl, 2 mM MgCl_2_, 0.1 mM EDTA, and 10% D_2_O. Error bars were determined via the covariance matrix in fitting R1 and R2 decay curves and represent standard error propagation of the spectral noise for hnNOE values. Residues without data for any sample are colored grey in the x-axis labels. Data were collected at 600 MHz and 298 K. **D.** Correlation between dense phase viscosity measured via microrheology and average ^15^N-R2 rates. For viscosity, the average and standard deviation of experimental replicates are shown. For R2, the average and standard deviation across residues plotted in (**B**) are shown. For All-R, the exponential decay and fit error from the ^1^H-1D is also shown (open symbol, see Methods and Supplementary Figure 9). The All-R R2 is likely an underestimate of the true average since data are only from the few most mobile residues.

The data support that the H3 tails remain mobile on the ps-ns timescale within nucleosome condensates, while the increased R2 values could reflect i) the increase in viscosity, ii) a shift in the equilibrium between intra- and inter-nucleosomal tail-DNA interactions toward inter-nucleosomal interactions within the dense phase, or iii) a combination of the two. Previous in-cell NMR studies have provided a framework for understanding these data. Similar to the dense phase of a phase-separated sample, the higher viscosity within a cell is expected to result in decreased R1 values and increased R2 values, but an exaggerated effect is observed experimentally for R2 (77, 78). These in-cell observations can be explained by transient interactions with large biomolecules or complexes (77, 78). The minimal differences in R1 that we observe between the light and dense phases of unmodified NCP suggest that the large differences in R2 reflect an increase in inter-nucleosomal interactions within the dense phase. This is consistent with the known role of tail-DNA interactions in nucleosome condensates (7, 16, 21). However, we cannot rule out the possibility that the large differences in R2 could be due to R_ex_ contributions on the ms timescale.

### H3 tail basic residues govern tail mobility within NCP condensates

We next investigated nucleosomal H3 tail mobility within a subset of the mutants toward the extremes of condensate changes in our NCP system: 4xR-Q, All-K, and All-R. ^1^H-^15^N spectral quality (in terms of sensitivity) was improved for All-K and 4xR-Q as compared to unmodified (**Supplementary Figure 6A-B**). Chemical shift differences (Δ*δ*) determined between the dense and dilute phases from ^1^H-^15^N HSQC spectra were equal to (for All-K) or smaller than (for 4xR-Q) those observed for unmodified (**Supplementary Figure 6C**). In contrast, spectral quality was poor for All-R, especially for the dense phase (**Supplementary Figure 6A-B**): in ^1^H-^15^N HSQC spectra, signal is largely only observed for the two TGG motifs and the extended ‘hinge’ region around the second TGG motif, the H3 tail regions known to be most dynamic (76). NMR analyses are reported below for All-R with the caveat that the signal quality of these samples is very poor, and the data should be treated as qualitative.

We repeated NMR spin relaxation experiments with the mutants to gain insight into their effect on ps-ns nucleosomal H3 tail dynamics. Within each phase of All-K and 4xR-Q, ^15^N-R1 values show a mild increase while ^15^N-R2 and hnNOE values decrease as compared to unmodified-NCP (**Figure 5A-C**, **Supplementary Figures 7-8**). A stronger effect is observed for 4xR-Q. The increased mobility of the H3 tail within 4xR-Q-H3-NCP was expected in the light phase due to a previous study under non-phase-separating conditions (59) and is consistent with arginine-DNA interactions constraining the mobility of the tails. The increased mobility of the All-K-H3 tail (albeit not as high as for 4xR-Q) is consistent with arginines forming stronger interactions with DNA than lysines. As with unmodified, differences in R1 and hnNOE between the phases are minor for All-K and 4xR-Q, and differences in dense phase ^15^N-R2 values are most pertinent for understanding changes in tail mobility within NCP condensates. As discussed above with unmodified-NCP, R2 profiles from the dense phase likely reflect changes in inter-nucleosomal interactions in addition to changes in viscosity. The stepwise decrease in dense phase R2 values when moving from unmodified to All-K to 4xR-Q (**Figure 5B**) could be a combination of the increased mobility of the tail (with weaker lysine-DNA interactions or arginine neutralization) and a decrease in inter-nucleosomal interactions (or a related change in the inter-nucleosomal interaction configuration). The minimal differences in R2 between the two phases of 4xR-Q could indicate that there are only minor differences in tail-DNA interaction configurations between the two phases. Minimal data are available for All-R, and while the data only report on the most dynamic residues, they hint at interesting trends for this mutant: i) hnNOEs from the light phase ‘hinge’ region are elevated and suggest that the high arginine content decreases ps-ns dynamics of the tail (**Figure 5C**), and ii) differences in R2 between the light and dense phases align with the trends observed for unmodified and All-K (**Figure 5B**, **Supplementary Figures 7-9**).

Together, these NMR data support mobility of the H3 tails on the ps-ns timescale within NCP condensates. Neutralizing arginines or substituting in a weaker DNA-interactor (i.e., lysine) increases the mobility of the H3 tails, while swapping arginine in for lysine has the opposite effect. The mutants tested via NMR show a correlation between average R2 values and viscosity (**Figure 5D**). Note that the average R2 for All-R is from only the most mobile residues and thus assumed to be an underestimate of the true average across the tail. Differences in R2 between the mutants and their two phases may report on differences in intra- vs. inter-nucleosomal interactions and the interaction network within condensates.

## DISCUSSION

Here, we show that nucleosome phase separation propensity and properties are influenced by the number, identity, and distribution of basic residues and their modifications within the H3 tail. We tested the effects of H3 tail arginine and lysine neutralization on the phase boundary and material properties of NCP condensates. Overall, neutralizations within the H3 tail decrease the phase separation propensity of the NCP and the corresponding viscosity of their condensates. Neutralizing arginines in the ‘terminal’ region of the tail has a stronger effect than in the ‘central’ region of the tail, supporting a positional dependence where residues that can reach further to bridge nucleosomes play a larger role in phase separation. In general, arginines within the H3 tail have a stronger contribution to phase separation than lysines. Shifting the H3 tail content towards lysines decreases condensate viscosity while shifting towards arginine is the one modification tested that leads to an increase in viscosity. The H3 tails remain highly mobile within nucleosome condensates unless arginine content is increased, and tail mobility correlates with condensate viscosity. We interpret our data in terms of tail-DNA interactions but acknowledge that we cannot rule out contributions from histone-histone interactions. The findings presented here highlight the intrinsic material properties of chromatin at the mesoscale. However, the transition from mesoscale condensates to microscale chromatin domains likely involves additional layers of regulation and spatial organization that merit future investigation. Together, our data support that modifying H3 tail basic residues has a differential direct effect on nucleosome phase separation (as a function of tail charge patterns), with implications for the direct effect of charge-modulating histone PTMs on chromatin organization via phase separation.

Here, the systematic perturbation of H3 tail basic residues begins to address the role of charge patterning in the histone tails in regulating chromatin phase separation. Charge patterning in IDRs is important in charge-driven phase separation (79) and is relevant within the nucleus. Charge blockiness of IDRs is important in controlling which transcriptional regulators get incorporated into MED1 Mediator condensates (80). A recent study investigated the charge patterning of the linker histone H1 C-terminal tail and its role in condensates formed with DNA (81). The C-terminal tails of linker histones are well-mixed sequences that are lysine-rich and generally arginine-poor. In the context of the H1 C-terminal tail, higher arginine as compared to lysine content increases phase separation propensity and decreases diffusion within the condensate (in terms of FRAP recovery). This is consistent with the higher viscosity of All-R-H3-NCP as compared to the unmodified-H3 sequence containing a mixture of lysine and arginine.

The H3 tails retain a high degree of mobility within NCP condensates. Modifying arginine residues via neutralization or swapping them out for a weaker DNA-interactor (i.e., lysine) has a larger effect than the condensate environment on ps-ns tail dynamics as observed via hnNOE. This highlights that the NCP condensate environment minimally perturbs tail mobility and also emphasizes the role of dynamics in the histone language (59, 68, 69). ^15^N-R2 values likely reflect differences in intra- vs. inter-nucleosomal interactions and network configuration within the condensate as modulated by basic residue modifications. Biological implications of retained tail mobility include that the tails are likely not occluded by the condensate environment from interacting with regulatory proteins in, for example, a transcriptionally active condensate. Perhaps other regulatory proteins within these nucleosome condensates are also highly mobile, but this remains to be tested. There is precedent for the high H3 tail mobility observed in NCP condensates. Within condensates of the H1 C-terminal tail with prothymosin-lZI, long-range chain reconfiguration of prothymosin-lZI is only slowed by a factor of three while the viscosity of the condensate is 300x greater as compared to the light phase (74). Additionally, the germ-granule protein Ddx4 forms a condensate that is ~50x more concentrated than the light phase but with similar hnNOE values (73). This contrasts with other systems that exhibit slowed dynamics in condensates. For example, hnRNPA2, the low-complexity domain of FUS, and an elastin-like polypeptide show larger changes in ^15^N-R1 and hnNOE values that are in line with a greater restriction in local motions (70–72).

Our results add to the emerging model of phase separation driving chromatin organization and regulation. Various nuclear factors are poised to tune condensate formation and physicochemical properties. It is anticipated that condensate viscosity influences the biological function of each chromatin subcompartment, with less viscous condensates associated with more active chromatin and vice versa (9, 14, 15). We propose that charge-neutralizing PTMs, such as lysine acetylation and arginine citrullination, tune phase separation to lower viscosity and allow for greater access to chromatin (**Figure 6**). The differences we observe in the phase separation propensity, condensate viscosity, and tail mobility within nucleosome condensates with charge-modulating mutations within the H3 tail support that the histone language (i.e., the positions and the types of modifications) mediates accessibility and biological function in part by tuning the biophysical properties of condensates. Furthermore, these data suggest that histone PTMs may largely tune chromatin phase separation in one direction, to lower viscosity, because many neutralize basic residues or add negative charge. In order to tune chromatin condensates to higher viscosity, additional contributions from linker DNA length or condensate-promoting proteins such as linker histone H1 and HP1lZI are necessary (7, 22, 23, 82, 83). Variations in histone sequence (histone variants) are also poised to influence phase separation, especially by altering the composition and relative prevalence of arginine and lysine (as seen for linker histone (81)). It remains an open question whether other interactions within chromatin influence phase separation (e.g., DNA sequence and methylation, the histone core, and tail-tail interactions). The environment and physicochemical properties of each phase-separated region of chromatin are anticipated to be determined by a combination of all nuclear factors present—chromatin, chromatin-interacting proteins, and epigenomic modifications.

**Figure 6.**
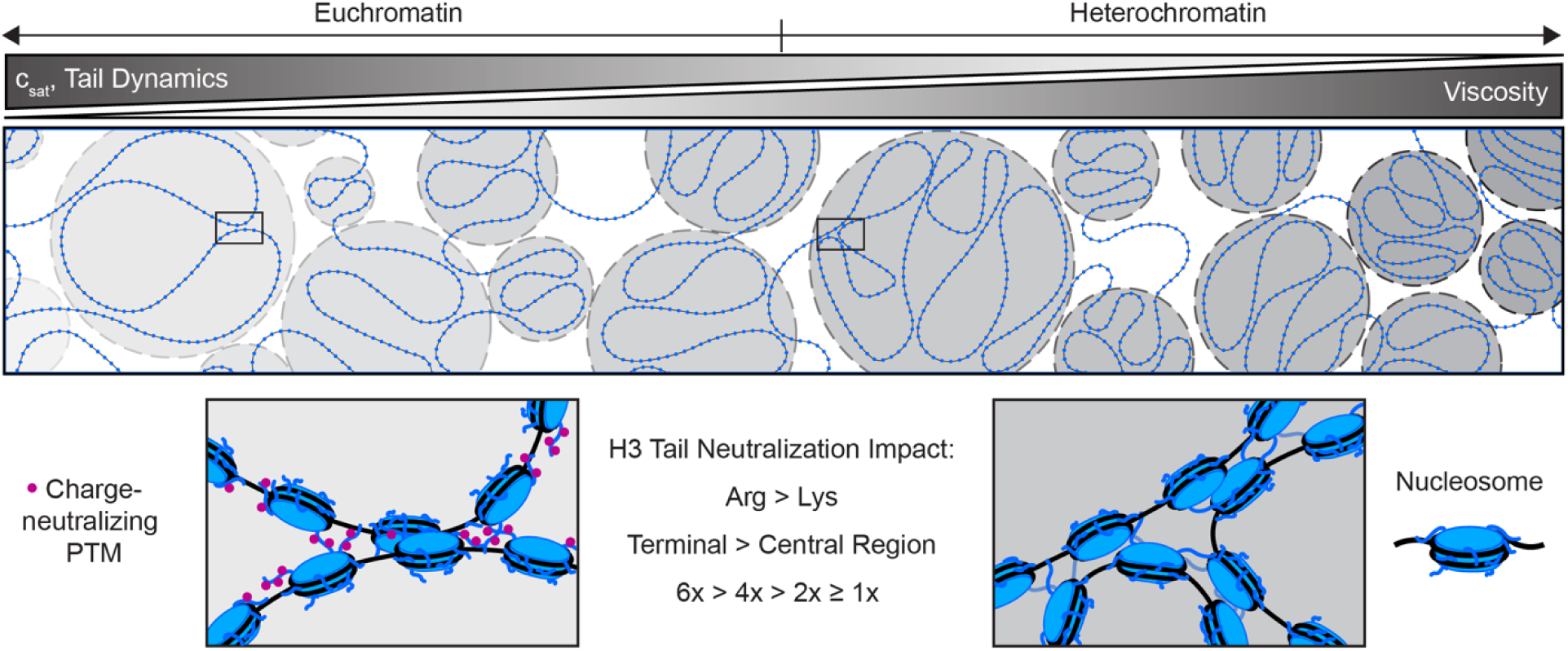
Histone tail charge patterning via modifications modulates chromatin condensates. In this model, charge-neutralizing PTMs, such as lysine acetylation and arginine citrullination, tune phase separation to lower viscosity and allow for greater access to chromatin. The differences observed in the viscosity and tail mobility of these nucleosome condensates suggest the histone language tunes accessibility and biological function in part by tuning the material properties of condensates.

## DATA AVAILABILITY

The data underlying this article are available in the manuscript, the supplementary materials, and upon reasonable request to the appropriate corresponding author. Unique plasmids are available from Addgene (plasmid # 240634) or upon request. The relaxation datasets presented in this study are deposited in the Biological Magnetic Resonance Data Bank (BMRB) with deposition numbers 53057, 53058, 53059, and 53458. All analysis and calculations for VPT were done using Python scripts from previous publications (54, 55) that are available on GitHub (https://github.com/BanerjeeLab-repertoire/Material-properties).

## SUPPLEMENTARY DATA

Supplementary data are available at *NAR* Online.

## ACKNOWLEDGMENTS

Thanks to Drs. Catherine Musselman, Karolin Luger, and Michael Poirier for gifts of histone plasmids.

## AUTHOR CONTRIBUTIONS

EM conceived of the initial project. EH, EM, DP, and PB designed the studies. EM, DP, and PB acquired funding and supervised the research. EH and SMZ prepared samples. EH, AS, KS, SY, RG, and EM performed the research and analyzed the data. EH, AS, KS, and EM visualized the data. EH and EM wrote the manuscript with contributions from AS, KS, DP, and PB.

## MATERIALS & CORRESPONDENCE

Correspondence on microrheology measurements should be addressed to Priya Banerjee at. Correspondence on simulations should be addressed to David Potoyan at. Other correspondence and material requests should be addressed to Emma Morrison at.

