## Supplementary Materials for "Histone H3 tail charge patterns govern nucleosome condensate formation and dynamics"

**Supplementary Methods:**

***Centrifugation assays.*** A centrifugation-based assay was used to determine the phase boundary of NCPs reconstituted with unmodified and mutant H3. Phase separation was induced at 300 µM NCP and room temperature (~23-24 °C) with a final volume of 30-60 µL. Final buffer conditions were 20 mM MOPS pH 7 (8 mM NaOH to pH), 0.1 mM EDTA, 150 mM KCl, and 2 mM $MgCl_2$. After allowing the sample to rest for 10 min, it was centrifuged at 18k rcf for 25 min at 25°C.

Following centrifugation, if phase separation occurred, the upper layer (light phase) was transferred to a separate tube, and both phases (now separate) were centrifuged again. Here, the concentration of the light phase was determined spectrophotometrically via NanoDrop absorbance measurement at 260 nm from three separate 1:10 dilutions in 2 M KCl. Using a Microman M25 positive displacement pipette (Gilson), 5 µL of dense phase (lower layer) was removed and transferred to 95 µL of MOPS buffer to dissipate phase separation. The concentration of this dense phase dilution was then determined by measuring absorbance at 260 nm from three separate 1:10 dilutions in 2 M KCl. Each plotted data point in Figure 1E is from one of these dilutions. If sufficient volume remained, a second dense phase measurement was obtained from this sample in the same manner, following a repeated centrifugation step. Statistical analyses were performed using R Statistical Software (v4.2.2; R Core Team 2022). A one-way ANOVA was initially used to determine variance across modified NCPs within either the light or dense phase measurements and was determined significant with a p-value of 5.37E-17 and 8.90E-59, respectively. To determine significance between modified NCPs (minimum n = 4 technical replicates and two biological replicates), a post-hoc Tukey's test was used. A p-value of $< 0.05$ was deemed significant for the pairwise comparisons between the NCP light phase and dense phase concentration measurements. Data were plotted using R Statistical Software (v4.2.2; R Core Team 2022).

If phase separation did not occur, the lack of two distinct layers following centrifugation was apparent. In this case, three separate 1:100 dilutions of the NCP sample were prepared in 2 M KCl, and the concentration was determined by measuring the absorbance at 260 nm. If the measured concentration matched the NCP input concentration (i.e., 300 µM), we concluded that the NCP does not phase separate under these experimental conditions. These measurements were excluded from the significance analysis due to the lack of light and dense phase concentration values.

**Supplementary Results:**

***Modulating H3 tail basic residues shifts the concentrations of the phase boundary.*** Centrifugation assays were used to measure the concentration of the light and dense phases ($c_L$ and $c_D$, respectively) at 25 °C (**Supplementary Figure 3C, Supplementary Tables 1-2**). These concentrations were measured at $c_L$ = 60 ± 20 µM and $c_D$ = 1000 ± 100 µM for unmodified-NCP. Tailless- and 6xK-Q-H3-NCP existed in only a single phase, and the measured concentrations (290 ± 30 µM and 300 ± 20 µM, respectively) matched the input concentration (300 µM) following centrifugation. The phase boundaries for 2xR-$Q_{terminal}$-H3-NCP were measured at $c_L$ = 100 ± 30 µM and $c_D$ = 960 ± 60 µM and for 2xR-$Q_{central}$-H3-NCP at $c_L$ = 70 ± 20 µM and $c_D$ = 980 ± 80 µM, supporting a shift in the onset of phase separation for neutralization of the terminal but not central region arginines. Phase boundary measurements support a narrowing of the boundary for 4xR-Q-H3-NCP as compared to unmodified-NCP, with the light phase concentration shifted to higher concentrations ($c_L$ = 90 ± 20 μM) and the dense phase concentration shifted to lower concentrations ($c_D$ = 750 ± 100 μM). By this approach, the additional neutralization of the two central arginines led to a lowering of the upper concentration boundary. The phase boundaries for 2xK-$Q_{terminal}$-H3-NCP were measured at $c_L$ = 80 ± 10 μM and $c_D$ = 1000 ± 100 μM, which are not significantly different from unmodified-NCP. While 4xK-$Q_{central}$-H3-NCP did not have a significantly shifted light phase concentration ($c_L$ = 80 ± 30 μM) as compared to unmodified-NCP, the dense phase was shifted to lower concentrations ($c_D$ = 670 ± 40 μM). Neutralization of the four arginine-adjacent lysine residues (4xK-$Q_{even}$-H3-NCP) resulted in a narrowing of the phase boundary, where light phase concentration shifted to higher concentrations ($c_L$ = 110 ± 10 μM) and the dense phase concentration shifted to lower concentrations ($c_D$ = 840 ± 40 μM) compared to unmodified-NCP. While this shifted boundary is qualitatively similar to the neutralization of the four arginines, neutralization of the four arginines has a larger effect on $c_D$, supporting a stronger effect of arginine neutralizations. The phase boundaries for All-K-H3-NCP were measured at $c_L$ = 42 ± 7 μM and $c_D$ = 1100 ± 50 μM, and for All-R-H3-NCP at $c_L$ = 50 ± 10 μM and $c_D$ = 950 ± 100 μM. Across these four concentration measurements, only the $c_D$ for All-K-H3-NCP was significantly different. All-K-H3-NCP is the only mutant tested with an elevated $c_D$ as compared to unmodified-NCP.

Systematic differences were observed throughout between the onset of droplets and turbidity (**Figure 2C-D**) and the $c_L$ measured after centrifugation of phase-separated samples. We speculate that these systematic differences could be due to small differences in temperature and the effect of centrifugal forces (i.e., non-equilibrium conditions). The dense- and light-phase concentration measurements carry larger uncertainties than ideal, in part due to the requirement for separating the phases by centrifugation, which introduces variability, and experimental error introduced by serial dilutions. For this reason, we place greater weight on brightfield microscopy and turbidity measurements, which probe phase behavior without mechanical perturbation. These assays consistently show a detectable and reproducible shift in the phase boundary upon neutralization of the N-terminal arginines.

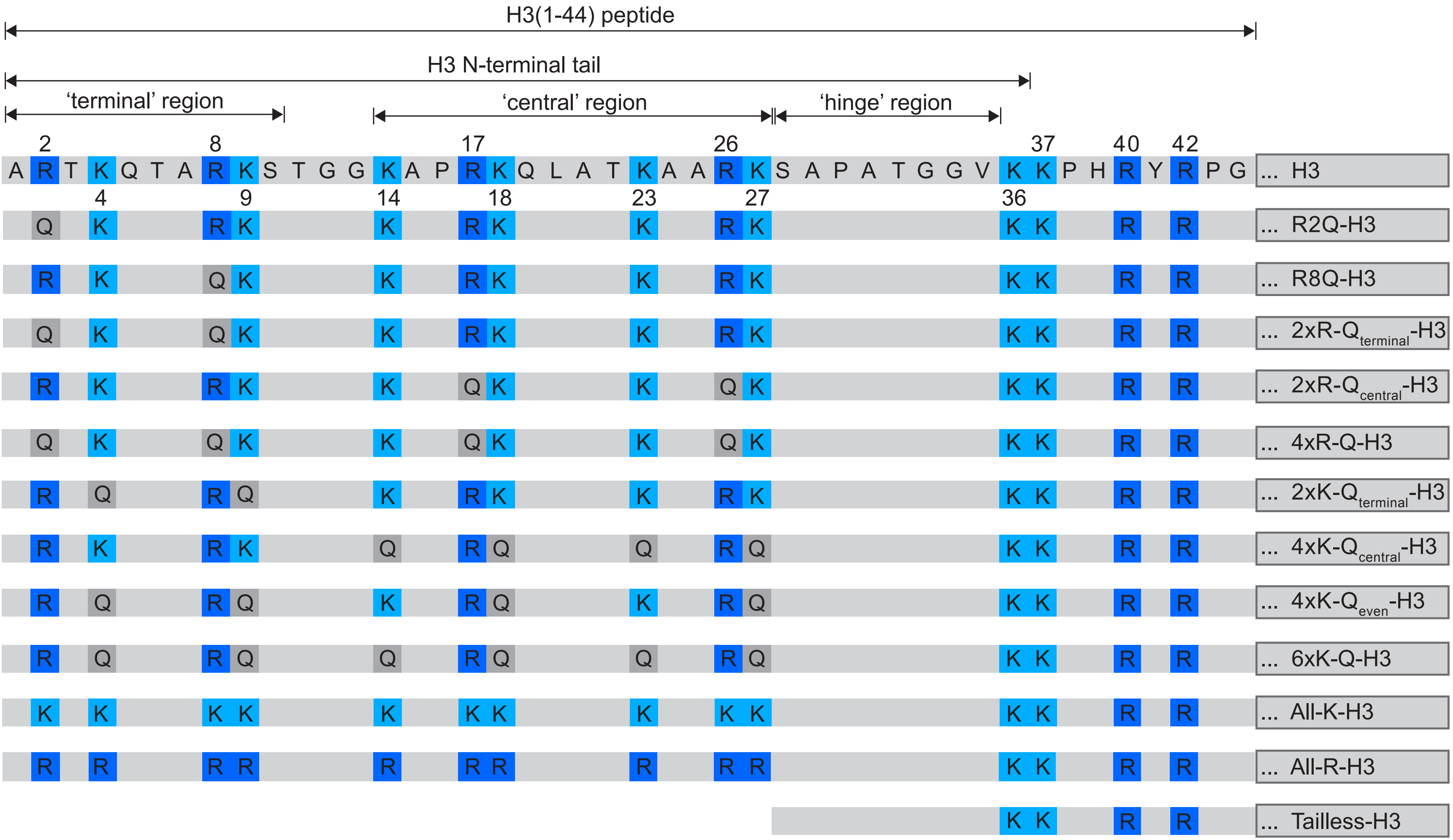


**Supplementary Figure1.** Histone H3 tail amino acid sequence that protrudes from the nucleosome core. Arginine (dark blue) and lysine (light blue) residues are highlighted and carried through to mutants below the unmodified-H3 sequence.

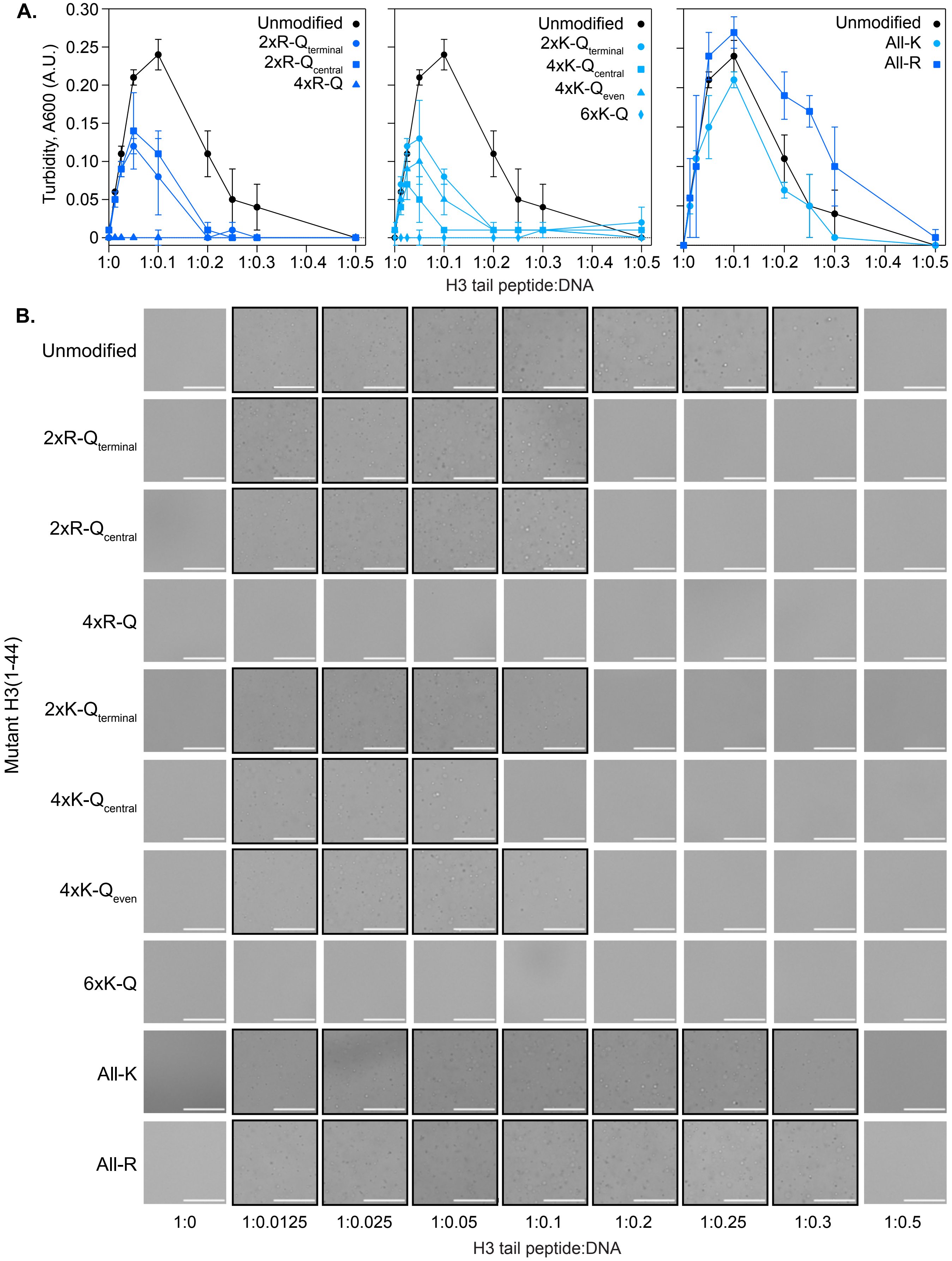


**Supplementary Figure 2. A.** Alternative plot of turbidity (absorbance at 600nm) where values (average of n=3) are shown as a function of H3 tail peptide:DNA ratio, corresponding to **Figure 2C**. Unmodified (black circles) H3 tail peptide is represented in each plot with mutant H3 peptides (blue shapes) where the arginine-to-glutamine mutants are plotted in the left panel, lysine-to-glutamine mutants in the middle panel, and the lysine-to-arginine and arginine-to-lysine mutants in the right panel. **B.** Brightfield microscopy images of unmodified and mutant H3 tail peptides collected at increasing H3 tail peptide: DNA ratios. All images are represented with a 50 μm scalebar (white). Images outlined in black indicate where droplets are observed.

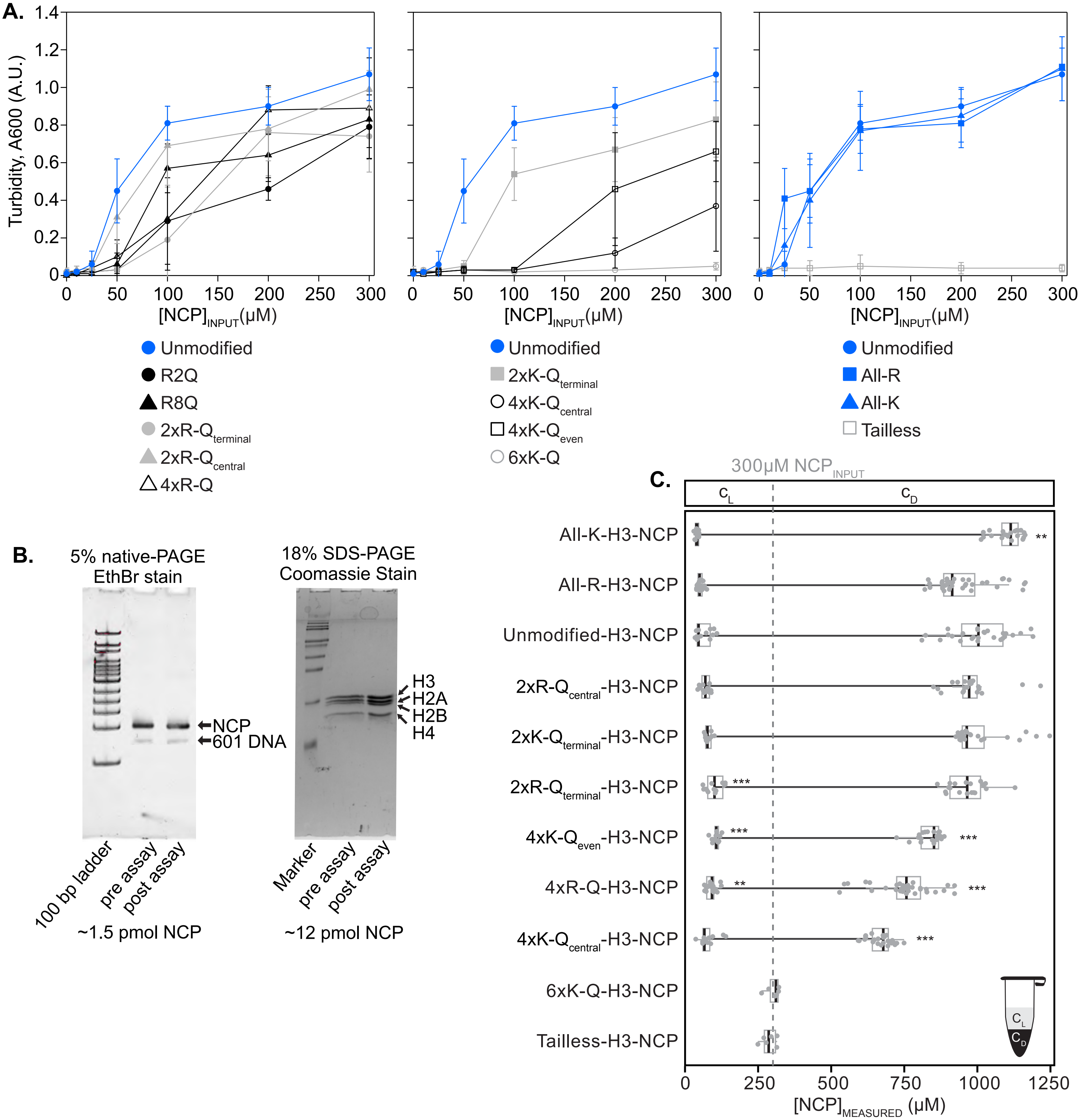


**Supplementary Figure 3. A.** Turbidity measurements for plate assays of unmodified (blue circles) and mutant (blue/grey/black) NCPs. Left plots show unmodified and arginine-to-glutamine mutant NCPs, center plots show unmodified and lysine-to-glutamine mutant NCPs, and right plots show unmodified, lysine-to-arginine, arginine-to-lysine, and tailless-H3-NCP. Alternative plots of turbidity (absorbance at 600 nm) values (average of n=3) are shown as a function of NCP concentration, corresponding to **Figure 2C**. **B.** Representative gel images used to assess NCP quality pre- and post-phase separation assay. NCP integrity represented in 5% native-PAGE and histone content represented in 18% SDS-PAGE. **C.** Phase boundary measurements for unmodified- and mutant-H3-NCPs following induction of phase separation at an input concentration of 300 μM NCP. The concentration of the light phase ($c_L$) and dense phase ($c_D$) were measured following centrifugation at 25°C. In the box plot, the black center vertical line marks the median, the grey box marks the interquartile range (IQR) from the 25th to 75th percentile, and the grey horizontal lines mark the minimum (25th percentile – 1.5*IQR) and maximum (75th percentile +1.5*IQR). A black line connects the $c_L$ and $c_D$ median values as a visual aid for the phase boundaries. Significance of $c_L$ and $c_D$ for each mutant as compared to unmodified-NCP was determined via ANOVA with Tukey post-hoc analysis where *** $p < 0.001$, ** $p < 0.01$, * $p < 0.05$. Those not marked are not considered significant.

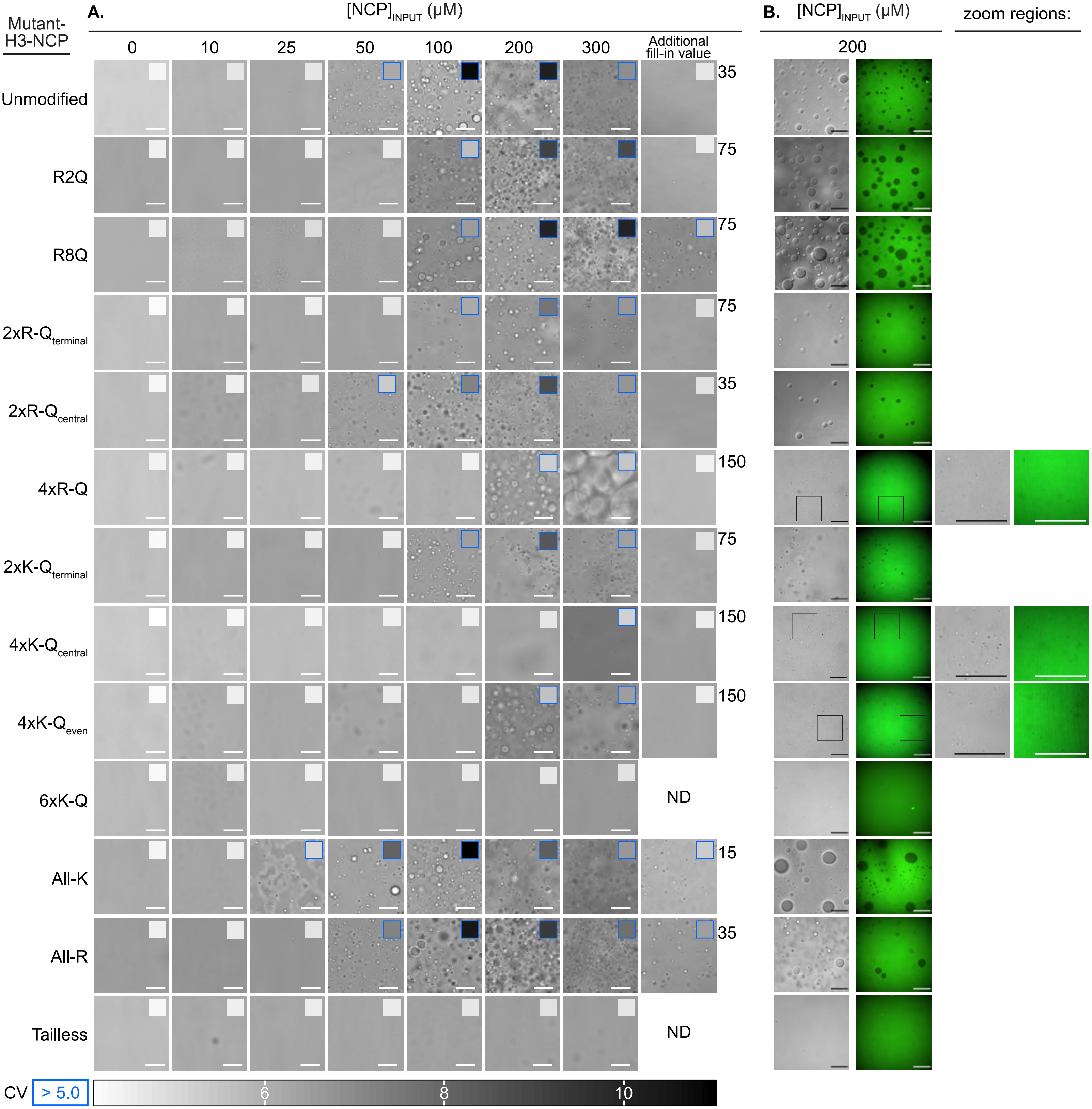


**Supplementary Figure 4. A.** Brightfield microscopy images of unmodified- and mutant-H3-NCPs collected at increasing concentrations of NCP. Each image is marked with a 50 μm scalebar (white, bottom right corner). Images for each concentration were collected with a minimum n = 3 and assigned a coefficient of variation value (see Methods), represented as a heatmap in the upper right corner of each image. Heatmap values greater than 5.0 are outlined in blue to indicate that droplets are observed. At 300 μM NCP, phase-separated samples of 4xR-Q-H3-NCP separated into two distinct layers by the time automated data collection began, falsely suggesting a lack of phase separation. For this mutant at 300 μM, an additional single-well experiment was collected to minimize the experimental dead time as much as possible. This single-well imaging is used in **Figure 1D** and is shown to the right of the image collected using the standard assay timing. **B.** Fluorescently stained unmodified- and mutant-H3-NCPs were collected at a concentration of 200 μM NCP, represented both as DIC (left image) and YOYO-1 fluorescence (right image) microscopy image with a 50 μm scalebar (bottom right corner).

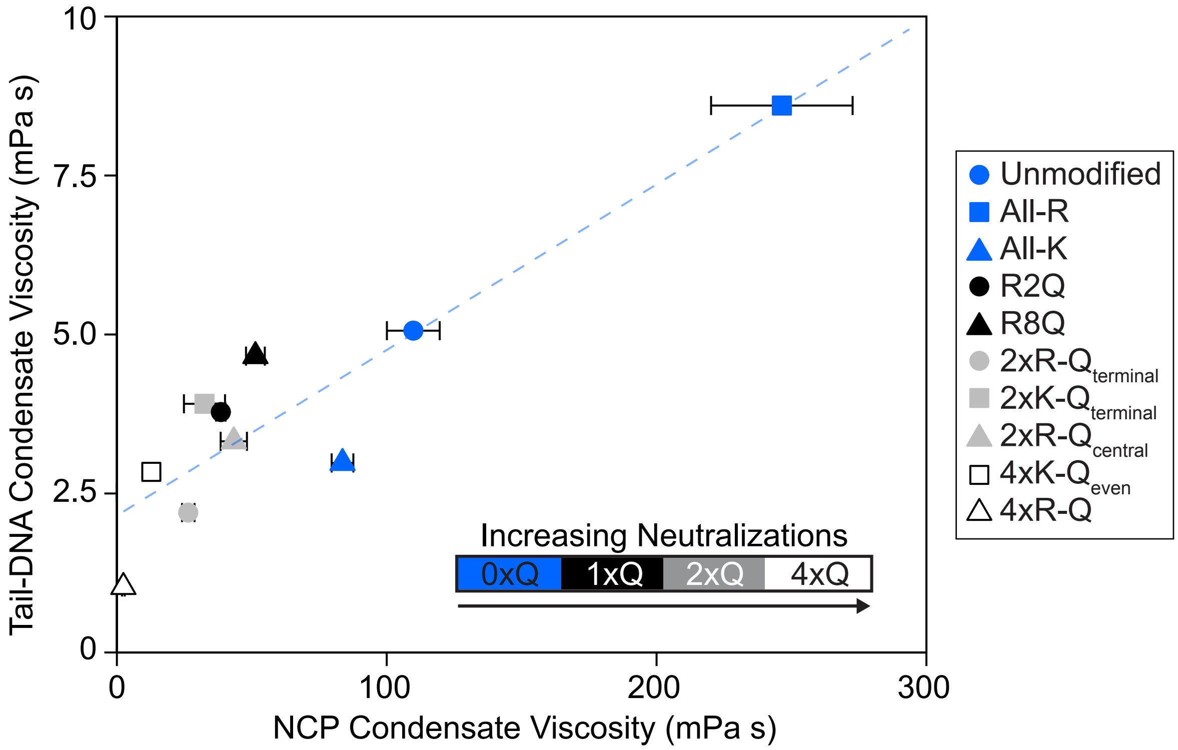


**Supplementary Figure 5.** Correlation between H3(1-44)-DNA condensate viscosity calculated from coarse-grained simulations and NCP condensate viscosity measured via microrheology. For NCP condensate viscosity, the average and standard deviation of experimental replicates are shown. Data symbols are consistent with **Figures 3 and 4**. The dashed blue line is a guide to the eye for the correlation.

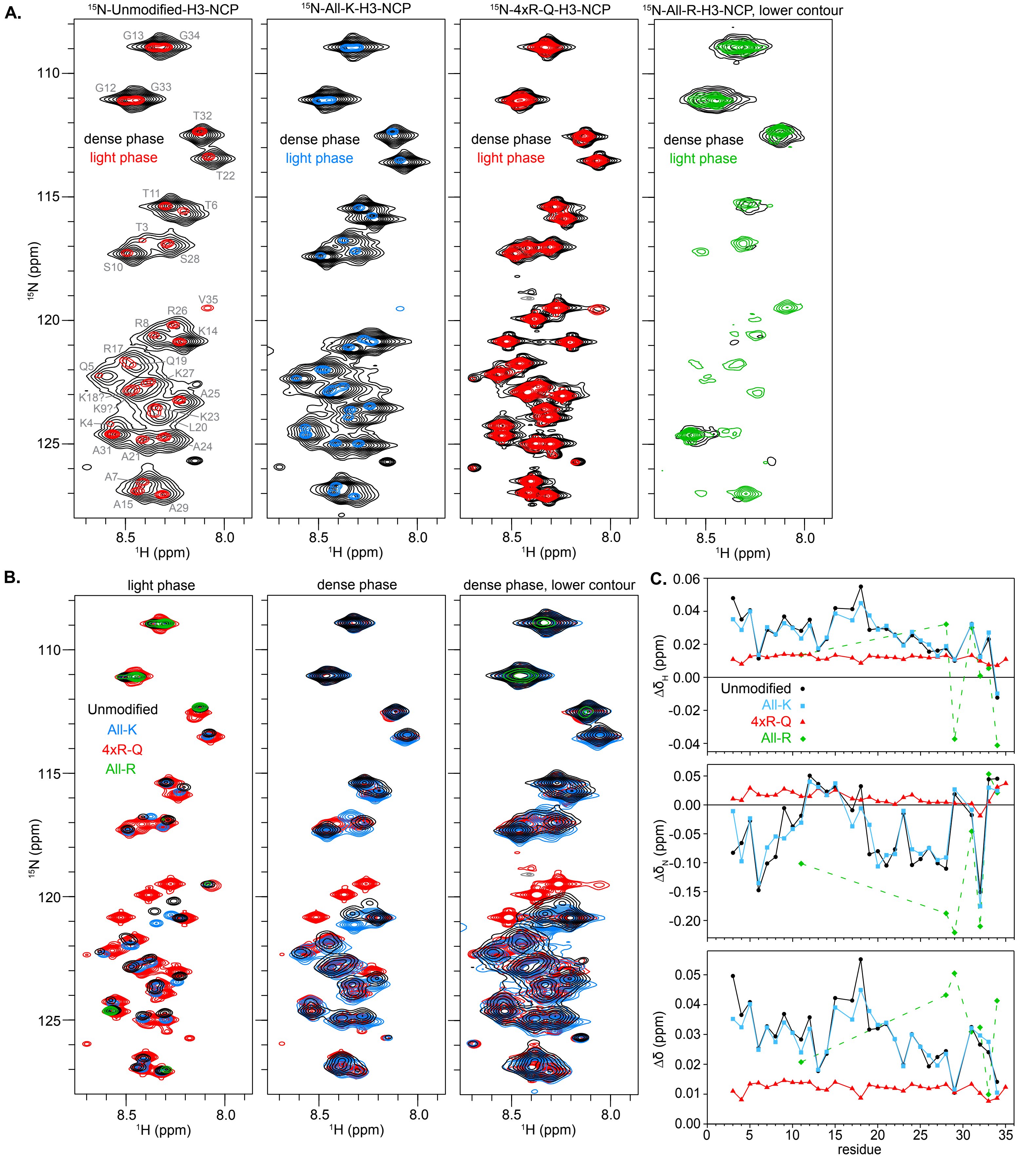


**Supplementary Figure 6.** Spectra of H3 tails within nucleosome condensates. **A.** Overlay of 1H-15N HSQC spectra of light (red, blue, or green) and dense (black) phases for each of 15N-Unmodified-, 4xR-Q-, All-K-, and All-R-H3-NCP. Spectra of the two phases are set to identical contours for a given NCP sample and are the same across the first three overlays; contours are lower for the fourth overlay. **B.** Overlay of 1H-15N HSQC spectra of all light and all dense phases separately for 15N-Unmodified-, 4xR-Q-, All-K-, and All-R-H3-NCP. Spectra are set to identical contours within a given overlay but are different between the overlays. **C.** Plots of chemical shift differences as a function of residue between light and dense phases for amide 1H ($\Delta\delta_H$) and amide 15N ($\Delta\delta_N$) separately and together ($\Delta\delta$). Data are shown for Unmodified- (black circles), All-K- (blue squares), 4xR-Q- (red triangles), and All-R- (green diamonds) H3-NCP.

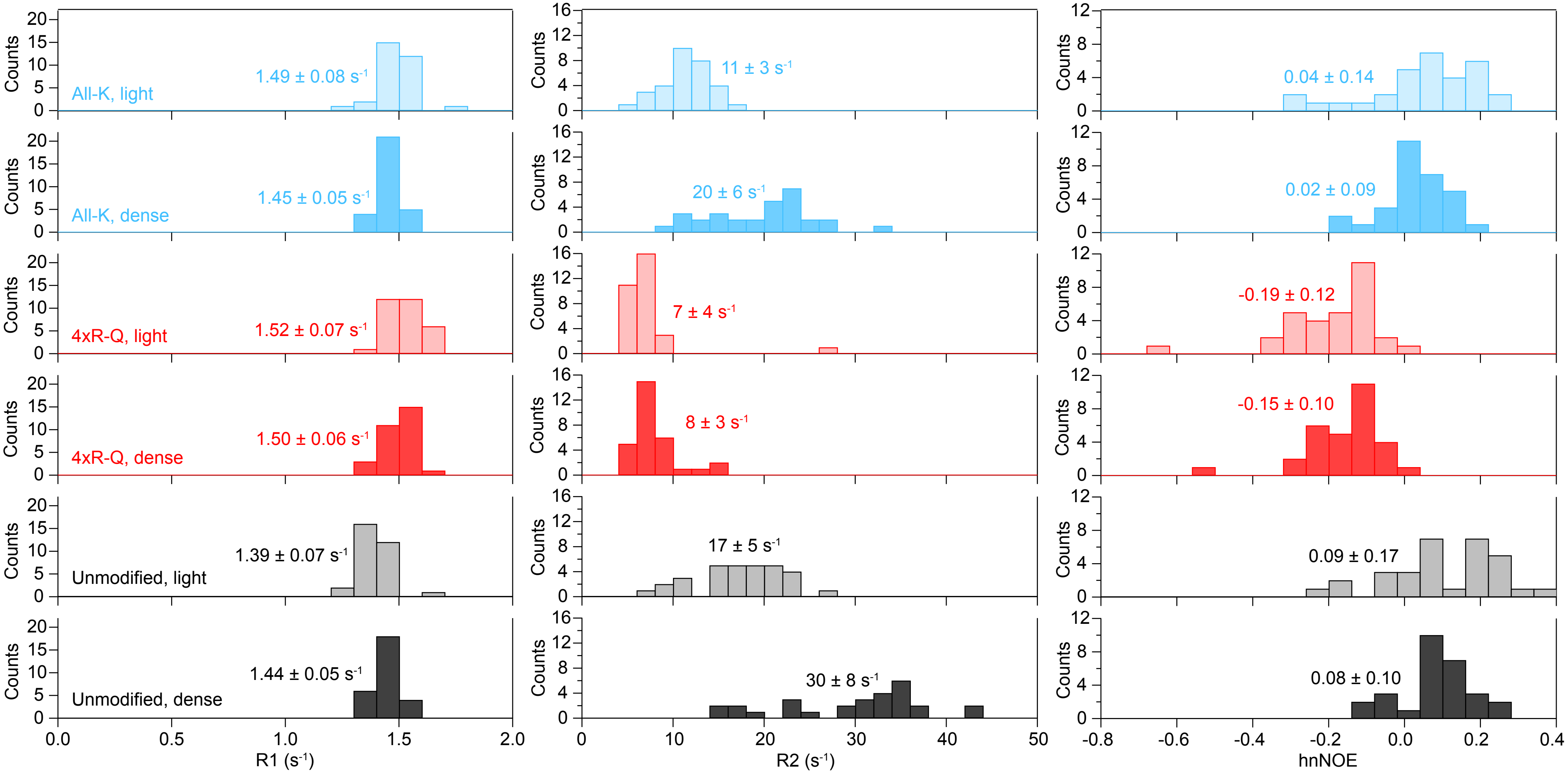


**Supplementary Figure 7. Histogram summary of NMR fast-timescale dynamics.** Histograms of 15N-R1 rates (left), 15N-R2 (central), and hnNOE (right) values for Unmodified- (black, lower plots), 4xR-Q- (red, central plots), and All-K- (blue, upper plots) H3-NCP with light and dense phases in lighter or darker shades, respectively. Averages with standard deviation are included on the plots.

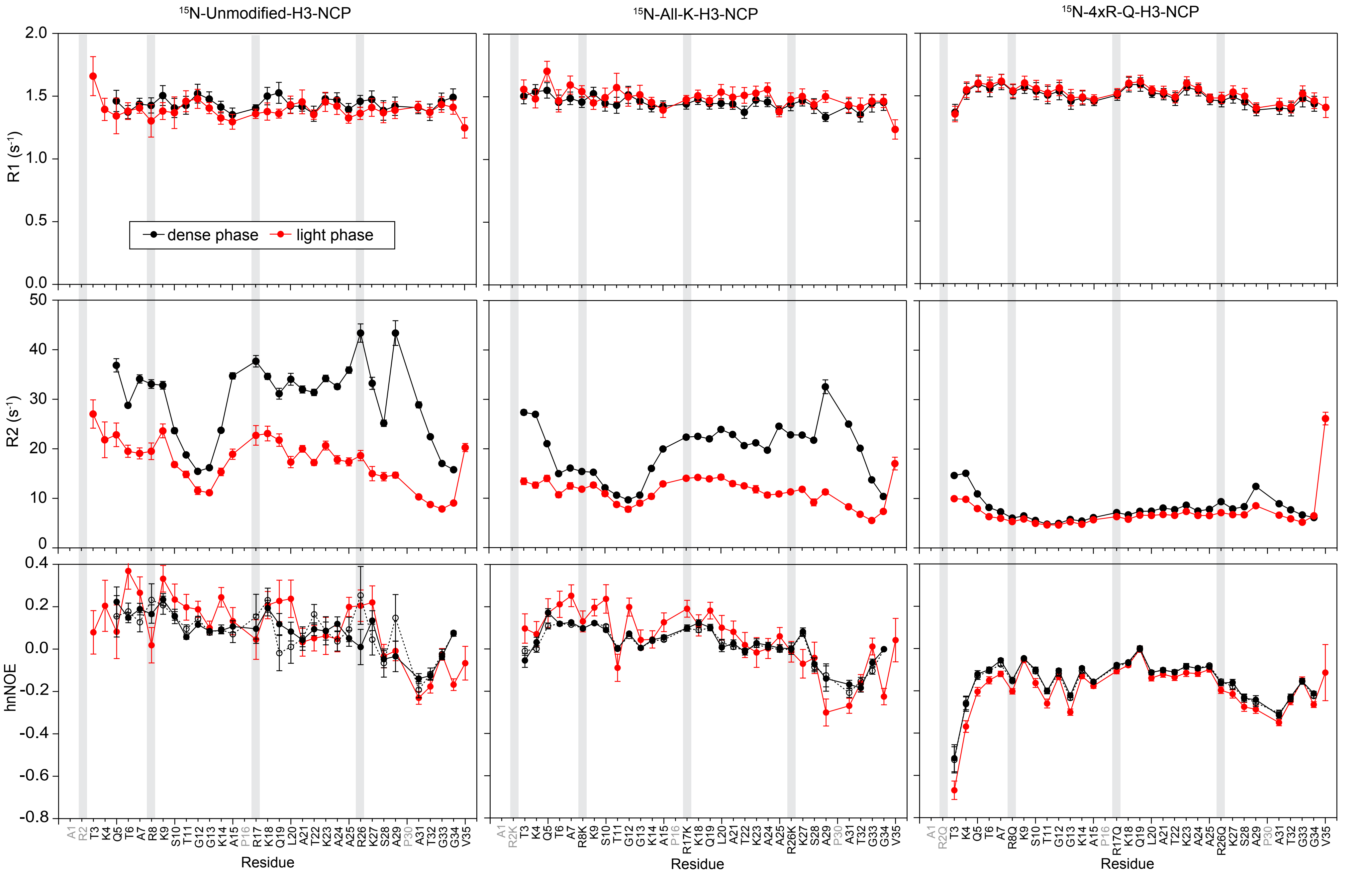


**Supplementary Figure 8. Summary of NMR fast-timescale dynamics.** Plots are shown for 15N-R1 rates (upper),15N-R2 rates (central), and hnNOE values (lower) as a function of residue for Unmodified- (left), All-K- (central), and 4xR-Q- (right) H3-NCP. Data were collected on the light phase (red) and dense phase (black) of NCP samples prepared in 20 mM MOPS pH 7, 150 mM KCl, 2 mM MgCl2, 0.1 mM EDTA, and 10% D2O. Mutated positions are highlighted by grey vertical bars to aid in visual comparison. Error bars were determined via the covariance matrix in fitting R1 and R2 decay curves and represent standard error propagation of the spectral noise for hnNOE values. Residues without data for any sample are colored grey in the x-axis labels. Data were collected at 600 MHz and 298 K.

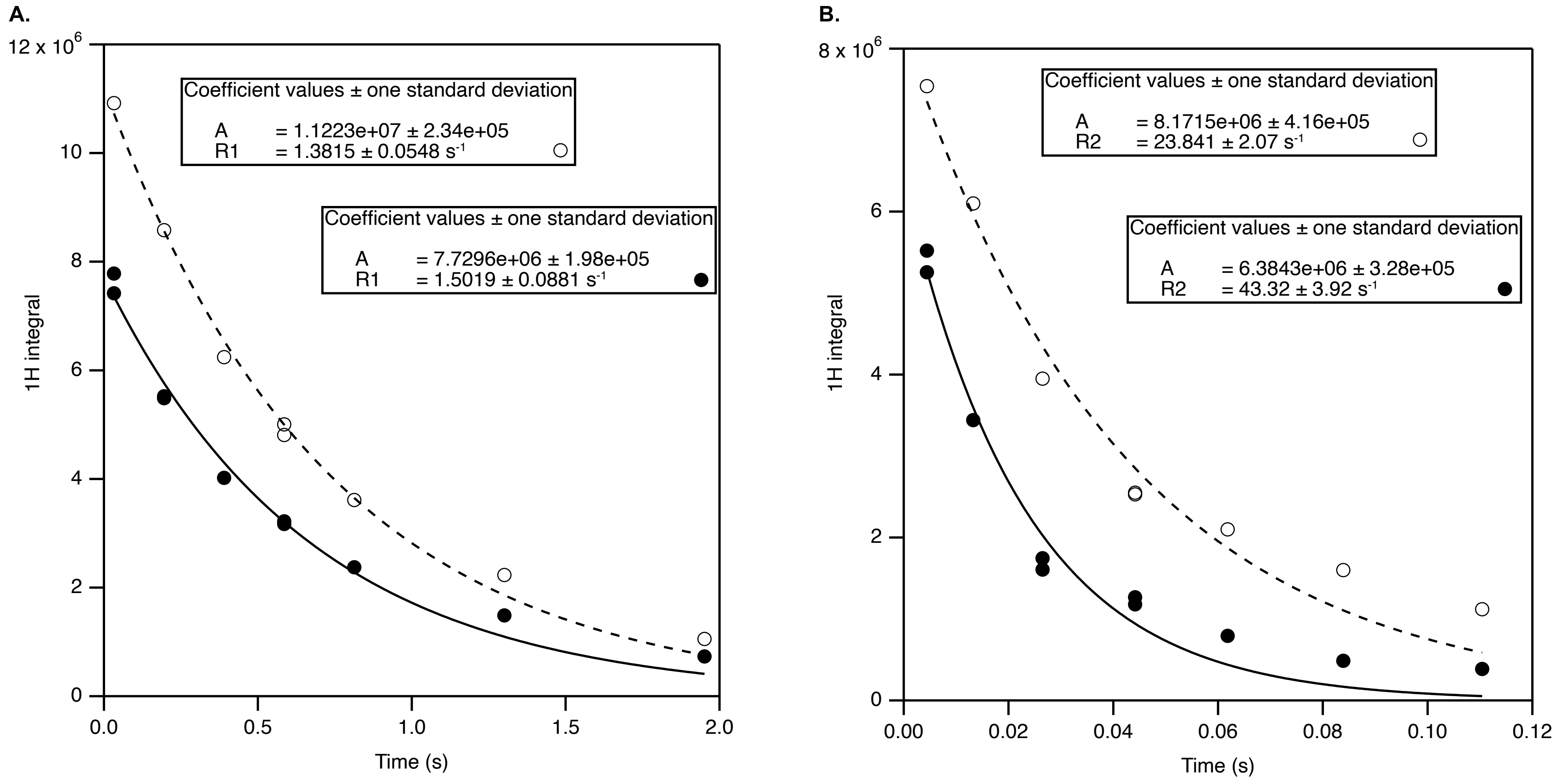


**Supplementary Figure 9. Relaxation decays from 1H integrals.** For 15N-All-R-H3-NCP, 1D versions of the R1 (**A**) and R2 (**B**) experiments were integrated over the amide region. Data were fit to a single-exponential decay without offset where the reported error is the estimated standard deviation of the fit coefficient. Fit values of R2 were plotted in Figure 5D (open green diamond). Signal is largely from the few most mobile residues and is thus not representative of the entire tail.

**Supplementary Table 1. Phase Boundary Measurements for Unmodified and Mutant H3-NCPs**

| NCP | $c_{L,\ measured}$ (µM) Ave | S.D. | $c_{D,\ measured}$ (µM) Ave | S.D. |
|---|---|---|---|---|
| Unmodifed-H3-NCP | 60 | 20 | 1000 | 100 |
| R2/8Q-H3-NCP | 100 | 30 | 960 | 60 |
| R17/26Q-H3-NCP | 70 | 20 | 980 | 60 |
| R2/8/17/26Q-H3-NCP | 90 | 20 | 750 | 100 |
| K4/9Q-H3-NCP | 80 | 10 | 1000 | 100 |
| K14/18/23/27Q-H3-NCP | 80 | 30 | 670 | 40 |
| K4/9/18/27Q-H3-NCP | 100 | 10 | 840 | 40 |
| R2/8/17/26K-H3-NCP | 42 | 7 | 1100 | 50 |
| K4/9/14/18/23/27R-H3-NCP | 50 | 10 | 950 | 100 |

| NCP | $[NCP]_{measured}$ (µM) Ave | S.D. |
|---|---|---|
| K4/9/14/18/23/27Q-H3-NCP | 300 | 20 |
| Tailless-H3-NCP | 290 | 30 |

Ave = Average   S.D. = Standard Deviation

**Supplementary Table 2. Phase Boundary Measurement Significance for Unmodified and Mutant H3-NCPs**

| NCP Light Phase Concentration ($c_L$) Comparison | p-value | |
|---|---|---|
| Unmodified-H3-NCP vs R2/8Q-H3-NCP | 2E-05 | *** |
| Unmodified-H3-NCP vs R2/8/17/26Q-H3-NCP | 1E-03 | ** |
| Unmodified-H3-NCP vs K4/9/18/27Q-H3-NCP | 6E-07 | *** |
| R2/8Q-H3-NCP vs R2/8/17/26K-H3-NCP | 2E-10 | *** |
| R2/8Q-H3-NCP vs R17/26Q-H3-NCP | 2E-03 | ** |
| R2/8Q-H3-NCP vs K4/9/14/18/23/27R-H3-NCP | 9E-09 | *** |
| R2/8Q-H3-NCP vs K14/18/23/27Q-H3-NCP | 4E-02 | * |
| R2/8/17/26Q-H3-NCP vs R2/8/17/26K-H3-NCP | 9E-09 | *** |
| R2/8/17/26Q-H3-NCP vs K4/9/14/18/23/27R-H3-NCP | 5E-07 | *** |
| R2/8/17/26K-H3-NCP vs R17/26Q-H3-NCP | 2E-02 | * |
| R2/8/17/26K-H3-NCP vs K4/9Q-H3-NCP | 1E-04 | *** |
| R2/8/17/26K-H3-NCP vs K4/9/18/27Q-H3-NCP | 5E-12 | *** |
| R2/8/17/26K-H3-NCP vs K14/18/23/27Q-H3-NCP | 1E-04 | *** |
| R17/26Q-H3-NCP vs K4/9/18/27Q-H3-NCP | 1E-04 | *** |
| K4/9Q H3-H3-NCP vs K4/9/18/27Q-H3-NCP | 2E-02 | * |
| K4/9Q H3-H3-NCP vs K4/9/14/18/23/27R-H3-NCP | 5E-03 | ** |
| K4/9/18/27Q-H3-NCP vs K4/9/14/18/23/27R-H3-NCP | 2E-10 | *** |
| K4/9/18/27Q-H3-NCP vs K14/18/23/27Q-H3-NCP | 3E-03 | ** |
| K4/9/14/18/23/27R-H3-NCP vs K14/18/23/27Q-H3-NCP | 7E-03 | ** |

| NCP Dense Phase Concentration ($c_D$) Comparison | p-value | |
|---|---|---|
| Unmodified-H3-NCP vs R2/8/17/26Q-H3-NCP | 0 | *** |
| Unmodified-H3-NCP vs R2/8/17/26K-H3-NCP | 4E-03 | ** |
| Unmodified-H3-NCP vs K4/9/18/27Q-H3-NCP | 9E-11 | *** |
| Unmodified-H3-NCP vs K14/18/23/27Q-H3-NCP | 0 | *** |
| R2/8Q-H3-NCP vs R2/8/17/26Q-H3-NCP | 4E-14 | *** |
| R2/8Q-H3-NCP vs R2/8/17/26K-H3-NCP | 3E-06 | *** |
| R2/8Q-H3-NCP vs K4/9/18/27Q-H3-NCP | 3E-05 | *** |
| R2/8Q-H3-NCP vs K14/18/23/27Q-H3-NCP | 0 | *** |
| R2/8/17/26Q-H3-NCP vs R2/8/17/26K-H3-NCP | 0 | *** |
| R2/8/17/26Q-H3-NCP vs R17/26Q-H3-NCP | 4E-14 | *** |
| R2/8/17/26Q-H3-NCP vs K4/9Q-H3-NCP | 0 | *** |
| R2/8/17/26Q-H3-NCP vs K4/9/18/27Q-H3-NCP | 9E-03 | ** |
| R2/8/17/26Q-H3-NCP vs K4/9/14/18/23/27R-H3-NCP | 3E-14 | *** |
| R2/8/17/26Q-H3-NCP vs K14/18/23/27Q-H3-NCP | 3E-03 | ** |
| R2/8/17/26K-H3-NCP vs R17/26Q-H3-NCP | 3E-05 | *** |
| R2/8/17/26K-H3-NCP vs K4/9Q-H3-NCP | 1E-02 | * |
| R2/8/17/26K-H3-NCP vs K4/9/18/27Q-H3-NCP | 2E-14 | *** |
| R2/8/17/26K-H3-NCP vs K4/9/14/18/23/27R-H3-NCP | 5E-09 | *** |
| R2/8/17/26K-H3-NCP vs K14/18/23/27Q-H3-NCP | 0 | *** |
| R17/26Q-H3-NCP vs K4/9/18/27Q-H3-NCP | 2E-06 | *** |
| R17/26Q-H3-NCP vs K14/18/23/27Q-H3-NCP | 0 | *** |
| K4/9Q-H3-NCP vs K4/9/18/27Q-H3-NCP | 7E-10 | *** |
| K4/9Q-H3-NCP vs K14/18/23/27Q-H3-NCP | 0 | *** |
| K4/9/18/27Q-H3-NCP vs K4/9/14/18/23/27R-H3-NCP | 1E-05 | *** |
| K4/9/18/27Q-H3-NCP vs K14/18/23/27Q-H3-NCP | 9E-10 | *** |
| K4/9/14/18/23/27R-H3-NCP vs K14/18/23/27Q-H3-NCP | 0 | *** |

Significance was determined via ANOVA with Tukey post-hoc analysis where *** $p < 0.001$, ** $p < 0.01$, * $p < 0.05$. Only significant comparisons are included in this table. Values are reported as 0 if less than 1E-14.

**Supplementary Table 3. Viscosities of NCP condensates measured at 24°C using VPT microrheology.**

| NCP | average viscosity (mPa s) | standard deviation (mPa s) |
|---|---|---|
| Unmodified-H3-NCP | 110 | 10 |
| R2Q-H3-NCP | 39 | 2 |
| R8Q-H3-NCP | 51 | 4 |
| R2/8Q-H3-NCP | 26 | 2 |
| R17/26Q-H3-NCP | 43 | 5 |
| R2/8/17/26Q-H3-NCP | 2.4 | 0.5 |
| K4/9Q-H3-NCP | 32 | 8 |
| K4/9/18/27Q-H3-NCP | 13 | 1 |
| R2/8/17/26K-H3-NCP | 84 | 4 |
| K4/9/14/18/23/27R-H3-NCP | 250 | 30 |

**Supplementary Table 4. Comparison of viscosities of NCP condensates measured at 24°C using VPT microrheology.**

| Condensate Viscosity Comparison | p-value | |
|---|---|---|
| Unmodified-H3-NCP vs R2Q-H3-NCP | 0 | *** |
| Unmodified-H3-NCP vs R8Q-H3-NCP | 0 | *** |
| Unmodified-H3-NCP vs R2/8Q-H3-NCP | 0 | *** |
| Unmodified-H3-NCP vs R17/26Q-H3-NCP | 0 | *** |
| Unmodified-H3-NCP vs R2/8/17/26Q-H3-NCP | 0 | *** |
| Unmodified-H3-NCP vs K4/9Q-H3-NCP | 0 | *** |
| Unmodified-H3-NCP vs K4/9/18/27Q-H3-NCP | 0 | *** |
| Unmodified-H3-NCP vs R2/8/17/26K-H3-NCP | 2E-5 | *** |
| Unmodified-H3-NCP vs K4/9/14/18/23/27R-H3-NCP | 0 | *** |
| R2Q-H3-NCP vs R2/8/17/26Q-H3-NCP | 2E-09 | *** |
| R2Q-H3-NCP vs K4/9/18/27Q-H3-NCP | 3E-05 | *** |
| R2Q-H3-NCP vs R2/8/17/26K | 0 | *** |
| R2Q-H3-NCP vs K4/9/14/18/23/27R-H3-NCP | 0 | *** |
| R8Q-H3-NCP vs R2/8Q-H3-NCP | 6E-05 | *** |
| R8Q-H3-NCP vs R2/8/17/26Q-H3-NCP | 0 | *** |
| R8Q-H3-NCP vs K4/9Q-H3-NCP | 6E-03 | ** |
| R8Q-H3-NCP vs K4/9/18/27Q-H3-NCP | 2E-10 | *** |
| R8Q-H3-NCP vs R2/8/17/26K-H3-NCP | 8E-08 | *** |
| R8Q-H3-NCP vs K4/9/14/18/23/27R-H3-NCP | 0 | *** |
| R2/8Q-H3-NCP vs R17/26Q-H3-NCP | 2E-02 | * |
| R2/8Q-H3-NCP vs R2/8/17/26Q-H3-NCP | 1E-04 | *** |
| R2/8Q-H3-NCP vs R2/8/17/26K-H3-NCP | 0 | *** |
| R2/8Q-H3-NCP vs K4/9/14/18/23/27R-H3-NCP | 0 | *** |
| R17/26Q-H3-NCP vs K4/9/18/27Q-H3-NCP | 4E-07 | *** |
| R17/26Q-H3-NCP vs K4/9/14/18/23/27R-H3-NCP | 0 | *** |
| R2/8/17/26Q-H3-NCP vs R17/26Q-H3-NCP | 1E-11 | *** |
| R2/8/17/26Q-H3-NCP vs K4/9Q-H3-NCP | 6E-07 | *** |
| R2/8/17/26Q-H3-NCP vs R2/8/17/26K-H3-NCP | 0 | *** |
| R2/8/17/26Q-H3-NCP vs K4/9/14/18/23/27R-H3-NCP | 0 | *** |
| K4/9Q-H3-NCP vs K4/9/18/27Q-H3-NCP | 3E-03 | ** |
| K4/9Q-H3-NCP vs K4/9/14/18/23/27R-H3-NCP | 0 | *** |
| K4/9/18/27Q-H3-NCP vs K4/9/14/18/23/27R-H3-NCP | 0 | *** |
| R2/8/17/26K-H3-NCP vs R17/26Q-H3-NCP | 3E-11 | *** |
| R2/8/17/26K-H3-NCP vs K4/9Q-H3-NCP | 0 | *** |
| R2/8/17/26K-H3-NCP vs K4/9/18/27Q-H3-NCP | 0 | *** |
| R2/8/17/26K-H3-NCP vs K4/9/14/18/23/27R-H3-NCP | 0 | *** |

Significance was determined via ANOVA with Tukey post-hoc analysis where *** $p < 0.001$, ** $p < 0.01$, * $p < 0.05$. Only significant comparisons are included in this table. Values are reported as 0 if less than 1E-14.

**Supplementary Table 5. Condensate parameters calculated from H3 tail-DNA coarse-grained simulations.**

| NCP | $c_{sat}$ (mg/mL) | viscosity (mPa s) |
|---|---|---|
| Unmodified-H3(1-44) | 10.5 | 5.1 |
| R2Q-H3(1-44) | 13.9 | 3.8 |
| R8Q-H3(1-44) | 11.6 | 4.7 |
| R2/8Q-H3(1-44) | 18.5 | 2.2 |
| R17/26Q-H3(1-44) | 14.2 | 3.3 |
| R2/8/17/26Q-H3(1-44) | 39.8 | 1.0 |
| K4/9Q-H3(1-44) | 12.2 | 3.9 |
| K14/18/23/27Q-H3(1-44) | 16.7 | 2.5 |
| K4/9/18/27Q-H3(1-44) | 19.6 | 2.8 |
| R2/8/17/26K-H3(1-44) | 18.3 | 3.0 |
| K4/9/14/18/23/27R-H3(1-44) | 6.0 | 8.6 |
