## Supplementary Video 1 Legend for "Histone H3 tail charge patterns govern nucleosome condensate formation and dynamics"

**LEGEND FOR SUPPLEMENTARY VIDEO**

**Supplementary Video 1.** Video moving through z-slices of microscopy images of unmodified-H3-NCP condensates with embedded 1 µm size yellow-green fluorescent carboxylate-modified beads. DAPI (blue), 200 nM in concentration, was added to the NCPs to show that the beads are embedded in the dense phase. Samples prepared identically but with the exclusion of DAPI were used for VPT microrheology.
